# Northern peatland microbial networks exhibit resilience to warming and acquire electron acceptor from soil organic matter

**DOI:** 10.1101/2024.07.17.603906

**Authors:** Katherine Duchesneau, Borja Aldeguer Riquelme, Caitlin Petro, Ghiwa Makke, Madison Green, Malak Tfaily, Rachel Wilson, Spencer W. Roth, Eric R. Johnston, Laurel A. Kluber, Christopher W. Schadt, Jeffrey P. Chanton, Paul J. Hanson, Susannah Tringe, Emily Eloe-Fadrosh, Tijana Del Rio, Konstantinos T. Konstantinidis, Joel E. Kostka

## Abstract

The microbial networks that regulate belowground carbon turnover and respond to climate change drivers in peatlands are poorly understood. Here, we leverage a whole ecosystem warming experiment to elucidate the key processes of terminal carbon decomposition and community responses to temperature rise. Our dataset of 697 metagenome-assembled genomes (MAGs) extends from surface (10 cm) to 2 m deep into the peat column, with only 3.7% of genomes overlapping with other well-studied peatlands. Unexpectedly, community composition has yet to show a significant response to warming after 3 years, suggesting that metabolically diverse soil microbial networks are resilient to climate change. Surprisingly, the dominant methanogens showed the potential for both acetoclastic and hydrogenotrophic methanogenesis. Nonetheless, the predominant pathways for anaerobic carbon decomposition include sulfate/sulfite reduction, denitrification, and acetogenesis, rather than methanogenesis based on gene abundances. Multi-omics data suggest that organic matter cleavage provides terminal electron acceptors, which together with methanogen metabolic flexibility, may explain peat microbiome resilience to warming.

## Introduction

Northern peatlands, which are often dominated by peat mosses (*Sphagnum* spp.; Walker et al., 2017) and characterized by their waterlogged organic soils (Rydin *et al*., 2013), contain approximately half of Earth’s soil carbon (Nichols and Peteet, 2019). The peatland “carbon bank” is a consequence of the intricate balance between plant productivity and slow decomposition rates of soil organic matter (SOM). *Sphagnum* spp. mosses are thought to play a key role in carbon sequestration. They act as ecosystem engineers, fostering acidic, nitrogen- and phosphate-poor conditions, and retaining water to create anaerobic environments that impede SOM decomposition (Van Breemen, 1995; Bragazza *et al*., 2013; Shao *et al*., 2023). Northern peatlands face mounting threats, with more pronounced warming projected at high latitudes in comparison to global averages (IPCC, 2013; Meredith *et al*., 2019; Masson-Delmotte *et al*., 2021; Taylor *et al*., 2022). The heterotrophic microbial networks that mediate SOM decomposition and the release of greenhouse gases (carbon dioxide, CO_2_; methane, CH_4_) are expected to be stimulated by warming (Treat *et al*., 2014; Gill *et al*., 2017; Hopple *et al*., 2020; Li *et al*., 2021; Wilson *et al*., 2021). However, our ability to predict the rates and controls of SOM decomposition in peatland soils is hindered by a limited understanding of the microbial networks involved.

Peatland soils differ profoundly in physical, chemical, and biological characteristics compared to their better understood upland forest counterparts. Upland forest soils are generally not saturated with water and oxygen is more available for microbial utilization. Thus, most SOM degradation is thought to occur under aerobic conditions and decomposition is hypothesized to be limited by carbon substrate availability (Kirk *et al*., 1976; Karhu *et al*., 2010). In contrast, water-saturated soils of *Sphagnum*-dominated peatlands are defined by anoxia, a paucity of inorganic terminal electron acceptors (TEAs), and respiratory pathways that mediate the terminal decomposition of organic matter are likely rate limiting (Duddleston *et al*., 2002; Hines *et al*., 2008; Drake *et al*., 2009; Bridgham *et al*., 2013). Carbon fixed by live vegetation is rapidly oxidized to CO_2_ above the water table, where oxygen is available (Duddleston *et al*., 2002; Hopple *et al*., 2020; Wilson *et al*., 2021). Immediately below the water table, fermentation products accumulate in peat porewaters in temperate and high latitude peatlands (Duddleston *et al*., 2002; Hopple *et al*., 2020), indicating that terminal electron accepting processes become rate limiting in the absence of oxygen. In this zone, SOM degradation is mediated by mainly anaerobic processes (Hopple *et al*., 2020; Wilson *et al*., 2021). Anaerobic laboratory incubation experiments corroborate field observations, showing that amendments of peat with energetically favorable, non-fermentable electron donors such as acetate and formate do not enhance mineralization, as evidenced by minimal changes in greenhouse gas production rates (Song *et al*., 2023). Conversely, a substantial increase in CO_2_ production is observed immediately after the introduction of nitrate (Song *et al*., 2023) and sulfate (AminiTabrizi *et al*., 2023), highlighting the limitation imposed on respiration by TEAs (Bodegom & Stams, 1999; Eriksson *et al*., 2010; Lozanovska *et al*., 2016; Song *et al*., 2023). Further, and despite TEA limitation under anaerobic conditions, several studies in temperate peatlands have consistently reported 10 to 1,000 times higher CO_2_ production relative to CH_4_ (Romanowicz *et al*., 1995; Chasar *et al*., 2000; Tfaily *et al*., 2014; Hodgkins *et al*., 2015; Wilson *et al*., 2016; Hopple *et al*., 2020; Song *et al*., 2023). Since the stoichiometry of methanogenesis predicts a 1:1 ratio of CO_2_ to CH_4_ production, higher ratios of CO_2_:CH_4_ implicate the predominance of alternative terminal electron accepting pathways (TEAP) (Conrad, 1999; Wilson et al., 2017). Fermentation products that accumulate at *in situ* temperatures are consumed at elevated temperatures, providing evidence that warming somewhat alleviates the limitation of terminal decomposition (Tveit *et al*., 2015; Song *et al*., 2023). This finding suggests that TEAs might be released from the degradation of organic matter itself, a hypothesis that remains to be tested.

A growing body of research has employed metagenomic approaches to uncover the community dynamics and metabolic potential of microorganisms in peat soils (Lin *et al*., 2014a; Lin *et al*., 2015; Woodcroft *et al*., 2018; St. James *et al*., 2021; Reji & Zhang, 2022; Richy *et al*., 2024). The focus has been on the initial steps in SOM degradation (carbohydrate-active enzymes) in acidic, carbon-rich peats dominated by *Sphagnum* spp., where members of the *Acidobacteriota* as well as microbial groups that mediate methane cycling were shown to dominate the soil microbial communities (Woodcroft et al., 2018; Wilson et al., 2021). Genes encoding diverse hydrolases believed to be specific for *Sphagnum*-derived carbon compounds were uncovered and the genomic potential for respiration of fermentation products (acetate) was linked to the *Acidobacteriota* at the oxic–anoxic interface (St. James *et al*., 2021). Extensive investigations at Stordalen Mire, a Swedish peatland containing a permafrost thaw gradient, led to the recovery of over a thousand metagenome-assembled genomes (MAGs) and the identification of *Methanoflorens* as a keystone group of methanogens in peatlands (McCalley *et al*., 2014; Mondav *et al*., 2014; Woodcroft *et al*., 2018; Ellenbogen *et al*., 2024). However, less attention has focused on terminal decomposition processes other than methanogenesis.

In this study, we examine the metabolic potential and microbe-metabolite interactions along a 2m depth gradient to uncover the main microbial networks that regulate belowground carbon turnover and leverage the SPRUCE experiment to begin elucidating their response to climate change drivers. The Spruce and Peatland Responses Under Changing Environments (SPRUCE) experiment, combines air and peat warming in a whole-ecosystem manipulation experiment which aims to understand the impact of climate change drivers (warming and elevated atmospheric CO_2_) on ecosystem functioning in a forested peat bog in northern Minnesota. During the first three years of the SPRUCE experiment, warming led to significant increases in greenhouse gas production (Hanson *et al*., 2020), and surface peat layers became more methanogenic as CH_4_ production rates increased more than CO_2_ production rates (Wilson *et al*., 2021). Preliminary metagenomic analysis suggested that, despite the increase in CH_4_ production, the abundance of methanogens remained stable (Wilson *et al*., 2021). Notwithstanding these recent insights, the influence of warming on other functional guilds, such as sulfate reducers or acetogens, has not yet been addressed, leaving a critical gap in our understanding of the effects of temperature on peat-dwelling microorganisms. In addition, the metabolic networks that drive the terminal steps of carbon turnover remain to be elucidated, hindering our ability to predict their response to climate change.

Our genome-resolved findings, supported by biogeochemical and metabolomics data collected in parallel, indicate that the keystone functional guilds of microorganisms mediating organic matter degradation in northern peat soils are resilient to warming. In the absence of oxygen, the functional potential for anaerobic respiration processes, such as sulfate-reduction and denitrification, exceeds the potential for methanogenesis. To circumvent the scarcity in inorganic TEA, peat microorganisms may obtain the TEAs used to fuel anaerobic respiration from the cleavage of organic matter. The wide metabolic versatility of abundant species may allow them to adapt their activity without significantly changing their abundance. For example, the dominant methanogens, Ca. *Methanoflorens* species, exhibit a genomic repertoire capable of performing both acetoclastic and hydrogenotrophic methanogenesis. The uncovered taxonomic and metabolic novelty of microbes along with our proposed conceptual model alter perceptions of anaerobic respiration pathways mediating terminal decomposition in peatlands and provide a mechanistic framework for improved predictions of the ratio of CO_2_:CH_4_ emissions.

## Results and Discussion

### SPRUCE peat harbors a plethora of novel taxa structured by depth

We sequenced 2.4 terabasepairs (Tbp) of metagenomic sequence data from 131 samples collected in 2015, 2016, and 2018 (5–40 Gbp of quality-processed data per sample), which are supported by biogeochemical and metabolomic data collected in parallel (**Supplementary Information** for a detailed description of the metabolic data). Metagenome binning resulted in the recovery of 697 medium- to high-quality MAGs (more than 50% complete and less than 10% contaminated, **Table S1**) dereplicated at the species level (i.e., average nucleotide identity [ANI] >95%). The MAG dataset well represents overall microbial richness at the site as evidenced by recruitment of the total metagenomic reads that could be mapped with representative MAGs. The lowest average read recruitment was observed at the peat surface 10-20 cm layer (28.7% of total reads) whereas at the three deeper depth intervals (40-50, 100-125, and 150-175 cm) on average 65%, 67.4% and 67.7% of total reads were represented in the MAG dataset, respectively (**Figure S1**). Depth was the main driver of variation in microbial community composition (P=0.002, adjR^2^=0.13, **Table 1, Figure 1A-B**) and functional potential (P<0.001, R^2^=0.623). Taxonomically, SPRUCE MAGs belong to a diverse range of bacterial (620 genomes) and archaeal phyla (77 genomes). Specifically, we recovered MAGs from 33 phyla (**Figure 1B-C**), including *Bacteria* belonging to Acidobacteriota (222 MAGs), *Actinomycetota* (75), *Proteobacteria* (64) and *Verrucomicrobiota* (48), and Archaea from the *Thermoproteota* (45) and *Halobacteriota* (17). Genomes of diverse aerobic chemoorganoheterotrophs were abundant in surface peat, while the deeper depths were predominated by taxa known for their anaerobic metabolism. *Acidobacteriota* was the most abundant phylum at all depths, with the highest abundances observed in surface peat (10-20 cm). Similarly, *Actinomycetota* and *Pseudomonadota* were more abundant in the surface layer. At mid depth (40-50 cm) we found a peak in the relative abundance of Desulfobacterota_B and *Halobacteriota* while the deep peat (100-125 and 150-175 cm) was the preferred habitat for *Thermoproteota*, *Thermoplasmatota*, *Verrucomicrobiota* and *Chloroflexota* (**Figure 1B**). This taxonomic stratification is a consequence of the metabolic distribution across the vertical profile in response to TEA availability and organic matter quality, as we show below.

**Figure 1.**
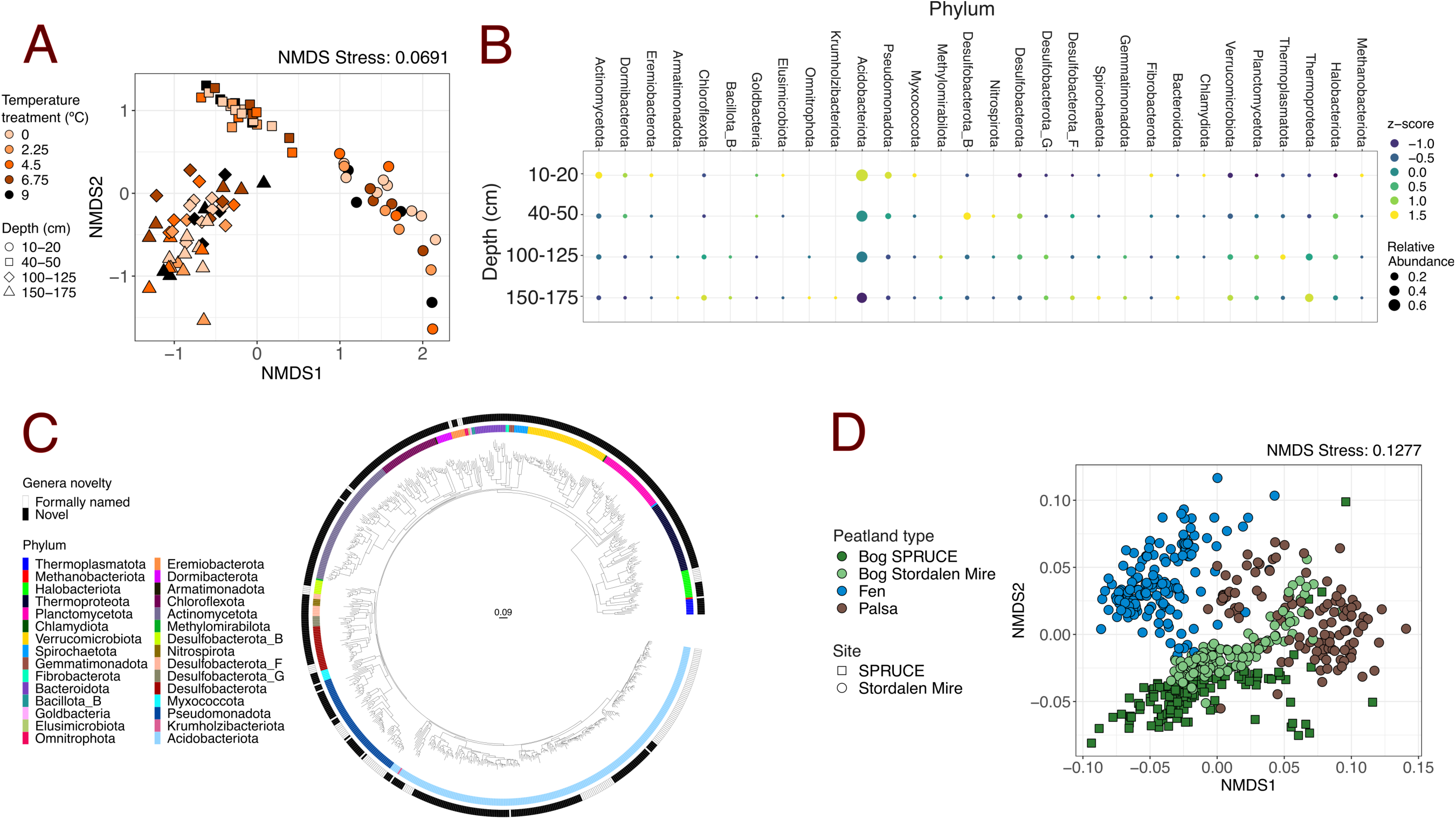
Taxonomic novelty and depth distribution of the metagenome-assembled genomes (MAGs) recovered from SPRUCE peat. A) Non-metric Multi-Dimensional Scaling (NMDS) plot based on MASH distances of metagenomes collected from SPRUCE in 2016 and 2018. The temperature treatment is indicated with colors while depth is shown with different shapes. B) Bubble plot showing the taxonomic distribution along the vertical profile. Dot size is proportional to the aggregated relative abundance (TAD80/GEQ) of each phylum while color shows the z-score value (the more positive the z-score is, the higher is the preference for that layer). Missing dots indicate the phylum was not detected at that depth. C) Phylogenomic tree based on 400 universal genes. Inner ring shows the taxonomic classification of each MAG at the phylum level. Outer ring indicates whether the MAG belongs to a formally named genera (white) or remains unnamed (black). Note that the majority lack a formal description. D) Non-metric Multi-Dimensional Scaling (NMDS) plot based on MASH distances of metagenomes from SPRUCE (squares) and Stordalen Mire (circles). Colors indicate the type of peatland (bog, palsa or fen). Note the clear geographical clustering of the samples.

The SPRUCE MAGs represent novel taxa, with the large majority (79.5%) belonging to genera that have not been formally described following the rules of either the ICNP or SeqCode. Furthermore, 20.8% and 4.6% of all MAGs were from an undescribed order and undescribed class, respectively (**Figure 1C**). The novelty of the SPRUCE soil microbiome was also supported by comparison against metagenomes retrieved from a broad range of global soil types (upland forest, tropical, Antarctic, grassland, and managed agricultural ecosystems) (**Figure S2**), revealing that SPRUCE peat harbors a plethora of ecosystem-specific, novel microbial taxa. To further examine the novelty of the dataset, we specifically compared our SPRUCE metagenomes against metagenomes from Stordalen Mire, a Swedish peatland with similar features to SPRUCE (type of soil, latitude, temperature, pH) for which the most detailed metagenomic analyses are publicly available (Woodcroft *et al*., 2018; Cronin *et al*., 2023). A non-metric multi-dimensional scaling (NMDS) plot based on MASH distances of raw reads revealed clustering by peatland type (bog, fen, palsa, P=0.001, adjR^2^=0.31, **Figure 1D, Figure S3**) and site (SPRUCE vs Stordalen Mire, P=0.001, adjR^2^=0.14, **Figure 1D, Figure S3**), further highlighting the novelty of these peat metagenomes with respect to previous work. In addition, only 26 of the total MAGs recovered in the present study (3.7%) shared more than 95% ANI to MAGs reported by Woodcroft *et al*. (2018) from Stordalen Mire. Most of these shared genomes were members of the phylum *Acidobacteriota* (8 MAGs), *Actinomycetota* (7 MAGs), and *Halobacteriota* (4 MAGs). Of particular interest, 5 out of the 26 MAGs belonged to Candidatus species named by Woodcroft *et al*. (2018). For example, SPRUCE 382 was assigned to Acidiflorens clade 2, SPRUCE 682 to *Ca*. Acidiflorens stordalenmirensis, SPRUCE 50 to *Ca*. Changshengia, SPRUCE 210 to *Ca*. Methanoflorens crillii, and SPRUCE 510 to *Ca*. Methanoflorens stordalenmirensis.

### The microbial community composition remains stable in response to warming despite changes in gas fluxes

The MAG composition-based NMDS and db-RDA analyses of samples collected in 2016 and 2018 did not reveal any significant effects of warming or eCO_2_ on overall microbial community composition at any of the sampled depths (**Figure 1A**). There were no significant changes in the relative abundance of any of the functional groups with temperature at any depth (**Figure 2**). Furthermore, the abundance of only a small subset of genomes (6.7%) appeared to correlate with the temperature treatments (**Table S2**). While the number of replicates per treatment may limit our statistical power to detect subtle changes, broadly the community demonstrates stability. Biogeochemical data from previous studies at SPRUCE (that is, using the same experimental design) have shown that microbial activity or function is significantly stimulated by elevated temperature. For example, the ratio of CO_2_ to CH_4_ in the peat declined with warming at the surface because of increased CH_4_ over CO_2_ production, indicating that the peatland ecosystem is becoming more methanogenic (Hopple *et al*., 2020; Wilson *et al*., 2021). Rate measurements of acetogenesis and methanogenesis conducted in summer of 2018 at SPRUCE, coinciding in space and time with our metagenomic dataset, also showed that rates of homoacetogenesis increased with temperature (LeeWays *et al*., 2022). Broadly, metabolite composition remained stable along the warming and eCO_2_ treatments through the depth profile (**Figure S4**). However, more thorough examinations of organic matter molecular composition at SPRUCE have revealed that the aerobic, surface layer of peat is degraded more readily with warming (Ofiti *et al*., 2022; Ofiti *et al*., 2023), possibly due to interactions between plants and their root associated microbes (Duchesneau *et al*., 2024). Thus, the response of peat microbes to warming at the community level must be (if any) smaller than the response in their activity. To identify potential mechanisms underlying the discrepancy between stable microbial community composition and unstable gas fluxes under warming, we performed a detailed study of the potential metabolic capabilities of peat microbes and developed a conceptual model. This model uses the depth profile as a proxy for carbon decomposition in peatlands, illustrating the microbial mechanisms driving CH_4_ and CO_2_ emissions, which can be tested for their activity later.

**Figure 2.**
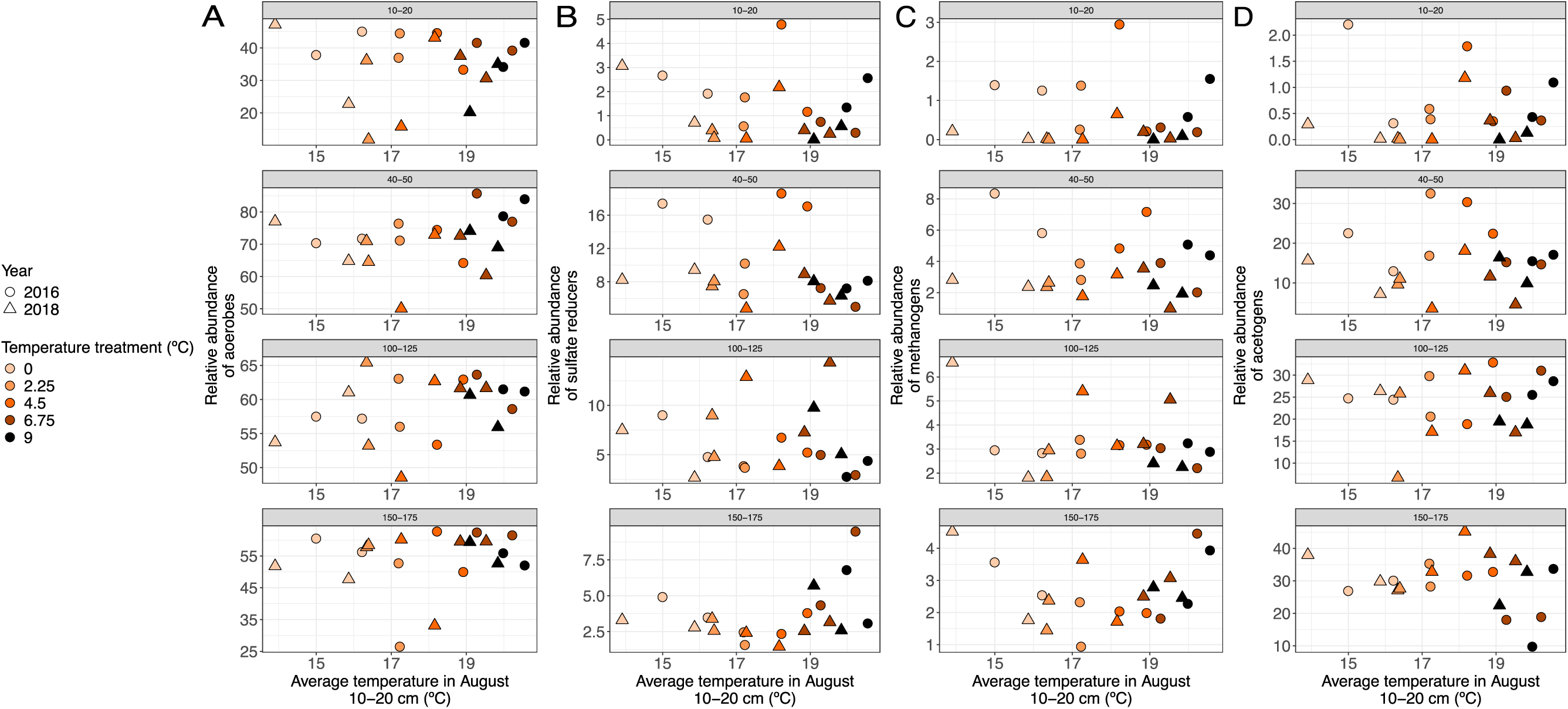
Warming treatment does not significantly impact the relative abundance of functional guilds. A) Mean aggregated relative abundance (TAD80/GEQ, x-axis) of MAGs encoding for aerobic respiration over the average temperature in August of the same year from 10-20 cm of the surface (°C, y axis). B) Mean aggregated relative abundance (TAD80/GEQ, x-axis) of MAGs encoding for sulfate reduction over the average temperature in August of the same year from 10-20 cm of the surface (°C, y axis). C) Mean aggregated relative abundance (TAD80/GEQ, x-axis) of MAGs encoding for methanogenesis over the average temperature in August of the same year from 10-20 cm of the surface (°C, y axis). D) Mean aggregated relative abundance (TAD80/GEQ, x-axis) of MAGs encoding for the Wood-Ljundahl pathway of acetogenesis over the average temperature in August of the same year from 10-20 cm of the surface (°C, y axis). Color indicates warming treatment, and shape indicates year when samples were collected. Each plot represents a distinct depth interval from 10 to 175 cm below the peat surface.

### Novel peat genomes reveal a broad metabolic diversity

We first focus on the most abundant MAGs, which remarkably exhibit a versatile metabolic potential, with the ability to perform two or more of the energy-generating pathways described above (**Figure S5**). This intricate interplay of microbial functionalities highlights the complex dynamics within peatland microbial communities. We present a few metabolic models before delving into our conceptual model to illustrate this metabolic flexibility.

A small but growing body of evidence reveals that members of the *Acidobacteriota*, a ubiquitous soil lineage, are capable of sulfate respiration in peat soils (Hausmann *et al*., 2018). Recent work based on lab incubations confirmed and extended these observations, showing the potential for both aerobic respiration and sulfate reduction in the same Acidobacteriota genome (Dyksma & Pester, 2023). *Acidobacteriota* MAGs in our dataset further corroborate these findings. For example, SPRUCE 682, with an average 3.2% relative abundance at the 40-50 cm, shows the potential to respire oxygen and/or sulfate. This genome, which belongs to the genus *Ca*. Acidiflorens (Sba1), also shows the potential to obtain sulfate by cleaving organic-sulfur compounds, such as choline-O-sulfate (**Figure 3**). SPRUCE 586 is another interesting genome within the *Acidobacteriota* phylum, increasing in abundance with depth and reaching 2.9% relative abundance in the 100-125 cm layer. Encoding the Wood-Ljungdahl pathway, SPRUCE 586 is a potential acetogen. Furthermore, *norB* was detected in this MAG, demonstrating its potential to produce N_2_O, a potent greenhouse gas (**Figure 3**). The family UBA7540, that contains SPRUCE 586, is among the most diverse families in the SPRUCE peat, with 17 species that are especially abundant in the deep peat (100-125 and 150-175 cm depths). However, only 35 UBA7540 genomes are found in GTDB and almost no information about this group is available in the literature. Thus, here we provide the first insights into the metabolic potential of this uncharacterized microbial group that abounds in the deep peat.

**Figure 3.**
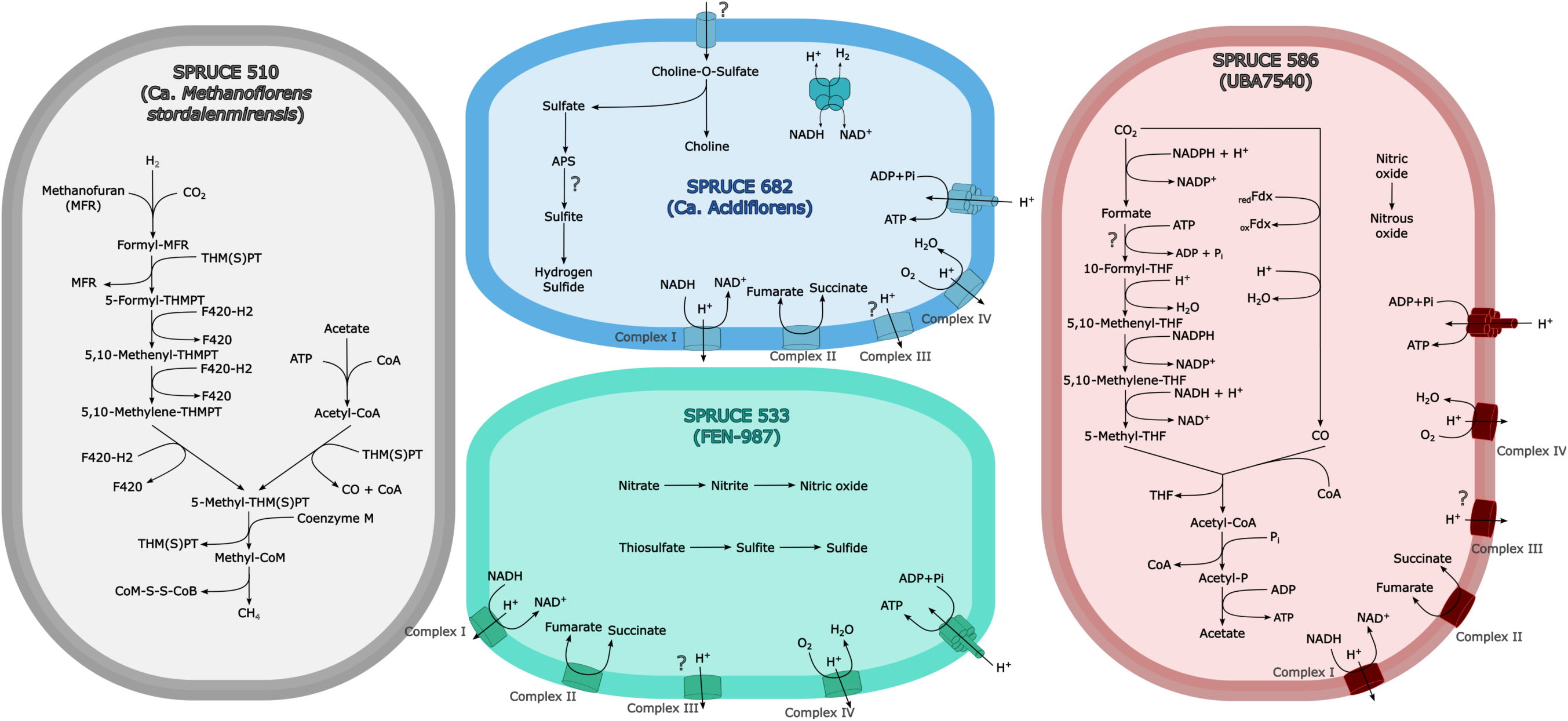
Versatility in metabolic potential for dominant species in SPRUCE peat. The metabolic potential of four abundant species is shown to highlight the capability of these species to adapt to the terminal electron acceptor availability. Question marks indicate enzymes or complexes that were not detected in the MAG but are likely to be present in the genome because the rest of the pathway in which they are involved was present.

The *Bathyarchaea* have been estimated to be among the most abundant soil microorganisms on Earth (Zhou *et al*., 2018). While more information is available from members of the phylum in marine sediments, the *Bathyarchaea* were detected in a range of anoxic sediments, including peatlands (Woodcroft et al., 2018). Here we show, for the first time, that the *Bathyarchaea* have the metabolic potential for sulfite reduction. Two MAGs belonging to the family FEN-987, SPRUCE 533 (0.07% mean relative abundance at 150-175 cm) and 422 (0.13% mean relative abundance at 100-125 cm), encode the corresponding hallmark genes. Interestingly, we observed the potential for oxygen, sulfite and nitrite respiration in SPRUCE 533, which highlights the adaptability of this species to variations in TEA availability (**Figure 3**).

Similar to other well-studied peatlands in Europe (Woodcroft *et al*., 2018), our previous work shows that genomes characterized as *Candidatus* Methanoflorens (Bog-38) are by far the most abundant methanogen genus present in SPRUCE soils (Wilson *et al*., 2021). The present study reveals that methanogens such as *Ca.* Methanoflorens may have a much more versatile metabolic potential than was previously perceived. For example, *Ca.* Methanoflorens stordalenmirensis (SPRUCE 510), the most abundant *Ca.* Methanoflorens species in our dataset (2.8% average relative abundance at 40-50 cm), encodes the potential for both acetoclastic and hydrogenotrophic methanogenesis (**Figure 3**). This dual potential was observed in 89.1% of the *Ca.* Methanoflorens MAGs (n=156) recovered from both SPRUCE and Stordalen Mire, indicating that it is not an exclusive trait of SPRUCE 510 or assembly/binning error (chimeric genome), but a general feature of this genus. To date, members of *Methanosarcina*, a genus that is largely considered as acetotrophic, are the only methanogens known to utilize both hydrogen and acetate as substrates to produce methane (Liu & Whitman, 2008). It should be mentioned, however, that experimental evidence that *Ca.* Methanoflorens can catalyze both acetoclastic and hydrogenotrophic methanogenesis remains to be confirmed.

### Conceptual framework for depth-stratified metabolic potential in a temperate northern peatland

The primary environmental forcing we observe is the distinct depth stratification of microbial community composition (**Figure 1A**) and functional potential (**Figure 4A**), which closely mirrors changes in redox conditions, gas emissions, and soil organic matter composition (**Figure 4B, S5, S4**). Redox chemistry drives microbial energy conservation pathways and is closely coupled to oxygen supply (Megonigal *et al*., 2004) linked to fluctuations in the water table, which rarely penetrates to below 30 cm depth at SPRUCE (Griffiths & Sebestyen, 2016; Hanson *et al*., 2020). This strong depth stratification of microbial communities and processes is in agreement with our past studies at the SPRUCE site (Lin *et al*., 2014b; Tfaily *et al*., 2014; Wilson *et al*., 2016; Tfaily *et al*., 2018), those of other peatland systems (Puglisi *et al*., 2014; Asemaninejad *et al*., 2019; Lamit *et al*., 2021) and as in our conceptual framework below, can be interpreted as a response of metabolic potential to changing terminal acceptor availability and soil organic matter quality.

**Figure 4.**
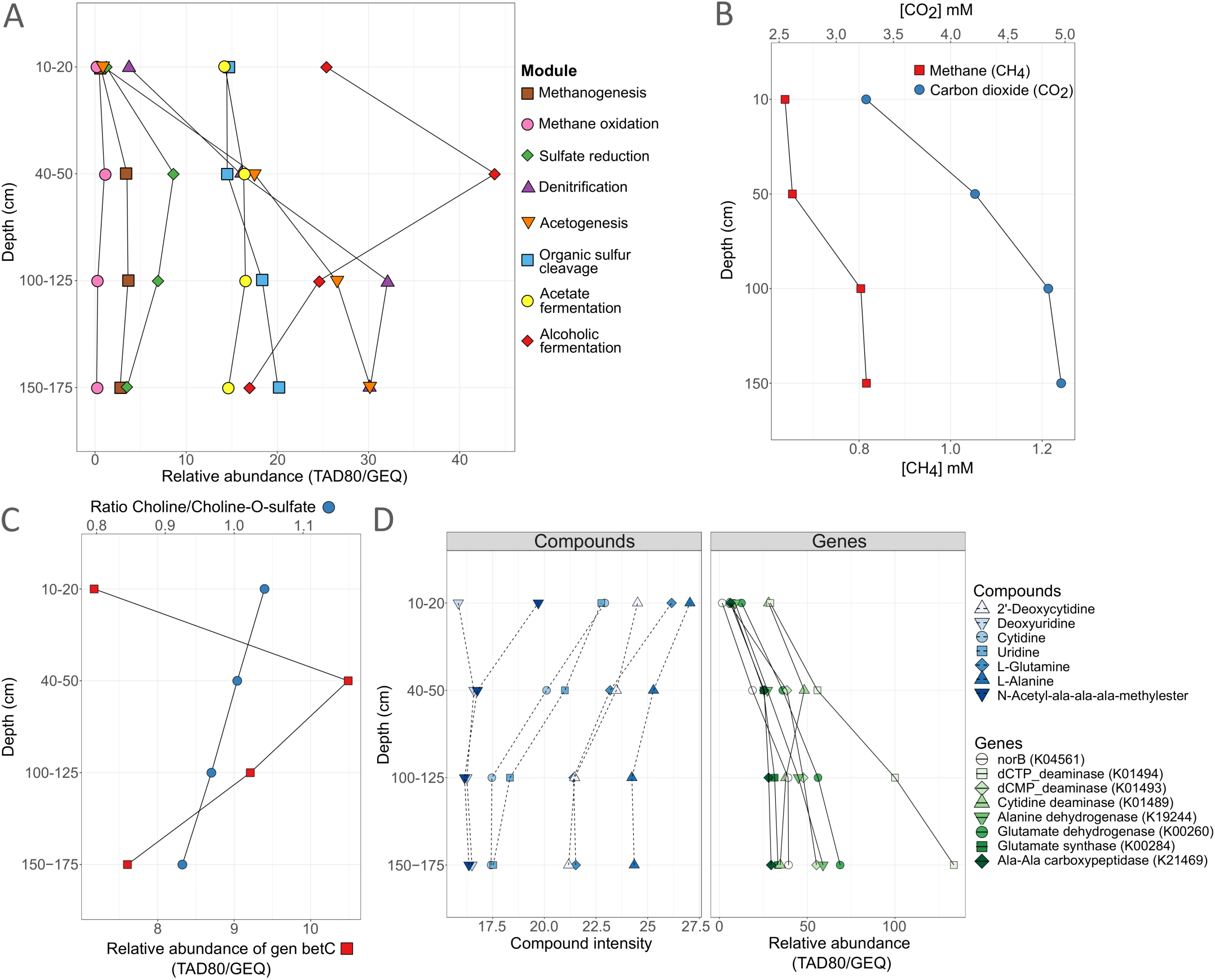
Data in support of the conceptual model of the terminal steps of organic matter degradation in peatlands. A) Mean aggregated relative abundance (TAD80/GEQ, x-axis) of MAGs encoding each metabolic pathway (module) along the depth profile (cm, y-axis). Each color and shape combination correspond to one metabolic pathway. B) Concentration of greenhouse gases CO_2_ (mM, top x-axis, blue circles) and CH_4_ (mM, bottom x-axis, red squares) along the peat column (cm, y-axis). C) Ratio of Choline to Choline-O-sulfate (intensity, top x-axis, blue circles) and the mean aggregated relative abundance (TAD80/GEQ, bottom x-axis, red squares), along the peat column (cm, y-axis). D) The intensity of compounds and the relative abundance of genes (TAD80/GEQ for genes and intensity for compounds, x-axis) related to deamination are depicted along the peat column (cm, y-axis). Each color and shape combination correspond to the depth profile of one gene or compound.

In SPRUCE peat, sulfate reduction is an important metabolic trait throughout the depth profile, detected in 34 MAGs identified as sulfate reducers across *Acidobacteriota* (11 MAGs), *Desulfobacterota* (including Desulfobacterota_B and Desulfobacterota_G, 13 MAGs), and *Thermoplasmatota* (2 MAGs, in the genus UBA184). Sulfite reduction, on the other hand, is attributed to 20 MAGs distributed among *Acidobacteriota* (9 MAGs), *Desulfobacterota* (6 MAGs), and *Thermoproteota* (2 MAGs in the family Fen-987). Within the *Acidobacteriota*, a substantial proportion, 5% of this taxonomic group, have the potential to mediate sulfate reduction, while 4% have the capability to perform sulfite reduction. Overall, the keystone functional guild of methanogens constitutes 3% of all MAGs, and notably, 20 MAGs within this group (n=21) exhibit the capability to execute multiple methanogenic pathways, further categorized into hydrogenotrophic (17 MAGs), acetoclastic (20 MAGs), and methylotrophic (6 MAGs) pathways (**Table S3**). The Wood-Ljungdahl pathway is common and phylogenetically diverse within our dataset; 58 bacterial MAGs, primarily from the phylum *Desulfobacterota* (15 MAGs), *Planctomycetota* (7 MAGs), *Acidobacteriota* (6 MAGs) and *Chloroflexota* (6 MAGs), show this potential. In addition, a large portion of the recovered *Thermoproteota* genomes contained the potential for homoacetogenesis (54.5% of the 33 identified MAGs). Note, however, that some species, such as sulfate-reducing bacteria, may use the reverse Wood-Ljungdahl pathway to generate energy through the oxidation of acetate to H_2_ and CO_2_ (Ragsdale & Pierce, 2008). Denitrification pathways are encoded by 174 MAGs, representing 25% of all MAGs, with Bacteria and Archaea contributing 160 and 14 of these MAGs, respectively. Members of the *Acidobacteriota* are significant players in denitrification, with 48 MAGs (21.6% of all *Acidobacteriota* MAGs, **Figure S6**) containing this potential, while *Desulfobacterota*, including Desulfobacterota_B and Desulfobacterota_G, contribute 23 MAGs, making up 46.7% of all *Desulfobacterota* involved in denitrification (**Figure S7)**.

### The acrotelm (10-20 cm): fluctuations in conditions lead to a diverse aerobic and fermentative microbial community

Peat mosses (*Sphagnum* spp.) carpet the soil surface in many temperate peatlands, including the S1 bog at SPRUCE sampled here. Beneath the living *Sphagnum* layer and above the water table, the 10-20 cm depth interval stands out as a hotspot for decomposition (**Figure S8**), as SOM derived from decaying plant biomass and labile organic compounds from root exudates are decomposed primarily via aerobic respiration. Our previous work showed that molecular oxygen is available at this depth (Lin *et al*., 2014b). Accordingly, MAGs encoding for pathways of aerobic respiration dominate at this depth (**Figure S8**). Fungi, which are either strictly aerobic or yeast capable of fermentation, have been also shown to peak in abundance (Lin *et al*., 2014b) and richness in this oxic/anoxic zone (Asemaninejad *et al*., 2017). We did not identify clear trends in the overall response of aerobes to warming **(Figure 2A**) and only a few aerobic genomes showed a positive (2 MAGs) and negative (4 MAGs) response to warming (**Table S2**), indicating a lack of response of this functional group to warming treatment.

The S1 bog water table occasionally reaches the 10 to 20 cm layer below surface and our previous work shows that oxygen is depleted where the soil is saturated with water (Lin *et al*., 2014b). Thus, the surface peat is ephemerally anoxic, and we find a high metabolic potential for acetate and alcoholic fermentation in the acrotelm (**Figure 4A, S8**), indicating heterotrophy is mainly driven by fermentation under anoxic conditions. In agreement with this observation, St James *et al*., (2020) detected the potential for acetate fermentation in most MAGs present in the 10-40 cm layer from the *Sphagnum*-dominated MacLean bog (New York State). In the present study, acetate production via fermentation is encoded mainly in the phyla *Verrucomicrobiota* (14 MAGs) and *Actinomycetota* (12 MAGs), while the potential for alcoholic fermentation was present in *Acidobacteriota* (165 MAGs), *Actinomycetota* (48 MAGs) and *Pseudomonadota* (46 MAGs). The total relative abundance of MAGs with the potential to ferment to acetate and alcohol did not respond to warming, and the response of MAGs capable of fermentation in the surface was mixed (**Table S2**).

In the acrotelm, aerobic respiration is coupled to canonical methanotrophy, at the oxic-anoxic interface where oxygen and methane are supplied from above and below, respectively. Anaerobic respiration pathways are less pronounced at this depth, with the genomic potential for methanogenesis, sulfate respiration, and acetogenesis at their lowest abundances (**Figure 4A)**.

### The mesotelm (40-50 cm): hot spot of anaerobic respiration via cleavage of TEAs from organic matter

Peatlands store the majority of their carbon in thick peat layers below the water table, where soil organic matter decomposition is dominated by anaerobic pathways (Megonigal *et al*., 2004; Drake *et al*., 2009; Bridgham *et al*., 2013; Wilson *et al*., 2016). Within the 40-50 cm depth range at SPRUCE, the potential for aerobic processes diminishes, giving their place to anaerobic respiration as the primary driver of organic matter degradation (**Figure 4A and S8**).

The paradigm in freshwater wetlands is that methanogenesis predominates over the TEA pathways coupled to the decomposition of SOM (Conrad, 1999; Megonigal *et al*., 2004; Drake *et al*., 2009; Bridgham *et al*., 2013; Tveit *et al*., 2015), thereby regulating the production of greenhouse gases (CO_2_, CH_4_). Here, however, despite a peak in methanogen abundance at 40-50 and 100-125 cm, the potential for other anaerobic respiration pathways (as reflected by relative abundance of the corresponding hallmark genes), such as sulfate reduction or denitrification, exceeds that of methanogenesis (**Figure 4A**). The stoichiometry of methanogenesis predicts a 1:1 ratio of CO_2_ to CH_4_ production and higher ratios of CO_2_:CH_4_ implicate alternate carbon oxidation pathways (Conrad, 1999; Wilson et al., 2017). Despite the scarcity of inorganic TEAs in peat, numerous field and incubation studies in northern peatlands, including those conducted at SPRUCE, have consistently reported 10 to 1,000 times higher CO_2_ production than CH_4_, indicating the dominance of alternative TEAPs in SOM decomposition (Romanowicz *et al*., 1995; Chasar *et al*., 2000; Tfaily *et al*., 2014; Hodgkins *et al*., 2015; Wilson *et al*., 2016; Hopple *et al*., 2020; Song *et al*., 2023). Our results provide further support for these observations (**Figure 4B**) but raise the question of the source of TEAs supporting anaerobic respiration, which are typically detected at very low concentrations in peat (Keller & Bridgham, 2007; Corbett *et al*., 2013). We propose that inorganic TEAs are obtained from the cleavage of the organic matter itself. Indeed, of the sulfate- and sulfite-reducers detected, 46 out of 55 MAGs demonstrate the genomic potential to cleave inorganic sulfur (sulfate, sulfite) from organic-sulfur compounds (**Figure S9**). This ability is widespread throughout the total microbial community, with 31% of our MAGs encoding genes for inorganic sulfur cleavage. Sulfur cleavage from organic matter is independent of oxygen regime (Lehnert et al., 2024) and persists throughout the depth profile (**Figure 4A**), suggesting that TEAs are cleaved from organic matter. Further, our data suggest that choline-O-sulfate, shown to be a major metabolite produced by *Sphagnum* (Carrell *et al*., 2022), is likely an important source of sulfate for anaerobic respiration, as the abundance of *betC* genes, responsible for cleaving SO_4_^2-^ from choline-O-sulfate, correlates with the ratio choline/choline-O-sulfate. Specifically, *betC* decreases in relative abundance in parallel with the abundance of detected choline-O-sulfate within our companion metabolite dataset and a relative increase in the byproduct, choline (**Figure 4C**). This suggests that bacteria capable of sulfate cleavage from organic matter might be utilizing it as an alternative source of TEA for respiration. Thus, our findings provide a possible explanation for the longstanding question of the mechanisms behind the unexpectedly high CO_2_ production in TEA-poor peatlands.

Our previous determinations of microbial activity corroborate the observations of genomic functional potential reported here, revealing a mid-depth hotspot in organic matter decomposition activity at 40-50 cm in the peat column, as evidenced by rates and ratios of greenhouse gas production, the accumulation of gases in soil porewaters, and the transformation of SOM (metabolomes) with depth (Lin *et al*., 2014a; Tfaily *et al*., 2014; Tfaily *et al*., 2018; Hopple *et al*., 2020; Wilson *et al*., 2021). While relative gene abundance provides compelling evidence for the importance of the metabolic potential of anaerobic respiration pathways at depth, our hypotheses must be confirmed by process-specific rate measurements in the future. Nevertheless, this potential mechanism of organic matter processing has implications for warming peatlands, and could explain increases in porewater CO_2_ observed below the water table (Wilson *et al*., 2021). Although warming did not impact the total relative abundance of MAGs with the potential to reduce sulfate, sulfite (**Figure 4B**), or cleave sulfate from organic matter, among the five genomes that significantly increase in relative abundance at 40-50 cm depth, two are *Acidobacteriota* involved in sulfur cycling (**Table S2**). The first, SPRUCE 490 in the family *Koribactericeae* (Coefficient=1.084, P=0.035), has the genomic potential to reduce sulfate and sulfite and ferment to alcohol. The second, SPRUCE 69, in the family *Bryobacteraceae* (Coefficient=1.650, P=0.45), can potentially cleave sulfate from choline-O-sulfate.

### The catotelm (100-125 and 150-175 cm): interactions between methanogens and acetogens and high potential for denitrification and methylotrophic methanogenesis

Below the mid-depth peak (40-50 cm) in anaerobic respiration activity, the genomic potential for acetogenesis via the Wood-Ljungdahl pathway continues to increase (**Figure 4A**). The pathways of methanogenesis are not partitioned equally throughout the depth profile, and although methanogens with the potential to produce methane via the acetoclastic and hydrogenotrophic pathway dominate throughout the peat profile, methylotrophic methanogens increase noticeably in the deep peat (**Figure S10**). Methylaminated organic compounds, such as 1-methyladenine and 1-methylguanine are negatively correlated (r=-0.84, and r=-0.84) with the abundance of the gene *mttB,* which encodes for a trimethylamine co-methyltransferase. The depletion of metabolites in the presence of key genes in the pathway of methylaminotrophic methanogenesis suggests that the pathway is active and links the deep peat to the complex, organic bound nitrogen cycle of peatlands. Of the few genomes that significantly respond to warming, SPRUCE 117 in the order *Methanomassiliicoccales*, which has the potential to produce methane using acetate, methanol, or methylamine as substrate, is negatively correlated with temperature in the deep peat (100-125 cm: Coefficient=-0.380, P=0.042, and 150-175cm: Coefficient=-2.206, P=0.020). Recently, methylotrophic methanogenesis has emerged as an important process in peatland soils, with reports of methylotrophic orders accounting for up to half of the transcription (Ellenbogen *et al*., 2024). In the bog where our experiment was conducted, ^13^C-labeled methylated substrates including methanol and monomethylamine were readily converted to CH_4_ by methylotrophic methanogens (Zalman *et al*., 2018).The loss of SPRUCE 117 with warming signals that methanogenic processes at SPRUCE are shifting with warming, but the decrease in metabolic potential for the demethylation of methylaminated organic compounds cannot explain why the bog is becoming more methanogenic with warming.

The deep peat exhibited significant metabolic potential for denitrification, with a peak occurring at 100-125 cm depth (**Figure 4A**). We observe a distinct distribution of genes involved in denitrification, with *norB* showing by far the highest relative abundance (**Figure S11**). This gene encodes nitric oxide reductase, an enzyme that catalyzes the reduction of nitric oxide (NO) to the potent greenhouse gas nitrous oxide (N_2_O), pointing to the catotelm as a potential source of N_2_O. Given the limited availability of oxidized nitrogen compounds in peat, the origin of the terminal electron acceptors for denitrification remains unclear. We propose that inorganic nitrogen forms could be derived from the organic matter itself, akin to sulfur compounds. Several lines of evidence from our metagenomic and metabolomic data support this hypothesis. Genes related to the hydrolysis and deamination of organic nitrogen compounds correlate positively with *norB*, while the respective compounds (e.g., nucleosides, peptides and amino acids) correlate negatively, suggesting consumption (**Figure 4D**). In agreement with this observation, we found peptidases encoded close to the *norB* genes (i.e., within 10 genes upstream or downstream) in 54% of all *norB*-encoding MAGs, which suggests that these genes may act on the same pathway. In addition, previous metabolomic studies at SPRUCE reported an enrichment of N compounds with depth (Tfaily *et al*., 2014). However, mineralized nitrogen is generally released in a reduced form as ammonium, and the potential for reoxidation or recycling of inorganic nitrogen in the deep, anoxic peat remains enigmatic. Possible explanations for the oxidation of ammonium in anaerobic peat soils include Fe(III)-mediated anaerobic ammonium oxidation (Feammox) (Tan *et al*., 2022; Wan *et al*., 2022), or ammonium oxidation by methanotrophs (Martikainen, 2022). While little information is available, biogeochemical evidence indicates Feammox is favored under acidic, anoxic conditions (Xan *et al*., 2022), which match the chemical environment in SPRUCE peat. Although Fe(III) concentrations at SPRUCE are low in comparison to mineral soils, a substantial pool of Fe(III) is observed that decreases with depth, reaching a minimum concentration at the 150–175 cm depth interval (Curtinrich *et al*., 2022). Indeed, we observed a strong correlation in the relative abundance of *norB* and ferritin genes (Pearson’s correlation=0.83; p-value=0), supporting a close relationship between them (**Figure S12**). Thus, these trends are consistent with our hypothesis that Fe(III) could be used as the electron acceptor in the oxidation of ammonium. Decomposition rates are shown to decline dramatically between 50 and 100 cm depth (Tfaily *et al*., 2014; Hopple *et al*., 2020), and an alternate explanation is simply that the cleavage of some oxidized organic S and N compounds supplies sufficient inorganic TEAs to support the lower rates of anaerobic respiration, and the replenishment of electron acceptor through reoxidation is not necessary.

### Concluding remarks

Our ability to predict the impact of climate change on the vast carbon stores of peatlands is hampered by a limited understanding of the microbial dynamics and metabolic pathways that regulate carbon turnover. Here, we contribute to filling this gap by providing a conceptual framework for the functional potential of microbial interaction networks extending 2 m into the peat column. The genome-resolved findings presented here provide an explanation for field observations that peatlands generally emit much more CO_2_ than CH_4_ under anaerobic conditions and propose mechanisms by which warming can impact the metabolism of the microbial community. Our conclusions on microbial dynamics in the belowground peat are supported by over 10 years of biogeochemical and metabolomics data collected from the SPRUCE site (Lin et al., 2014a; Lin et al., 2014b; Tfaily et al., 2018; Hopple et al., 2020; Wilson et al., 2021; Curtinrich et al., 2022; Petro et al., 2023; Song et al., 2023). From our conceptual framework, we describe two potential mechanisms that could explain why the microbial community maintains resilience despite increasing greenhouse gas emissions in the face of warming temperatures (**Figure 5**).

**Figure 5.**
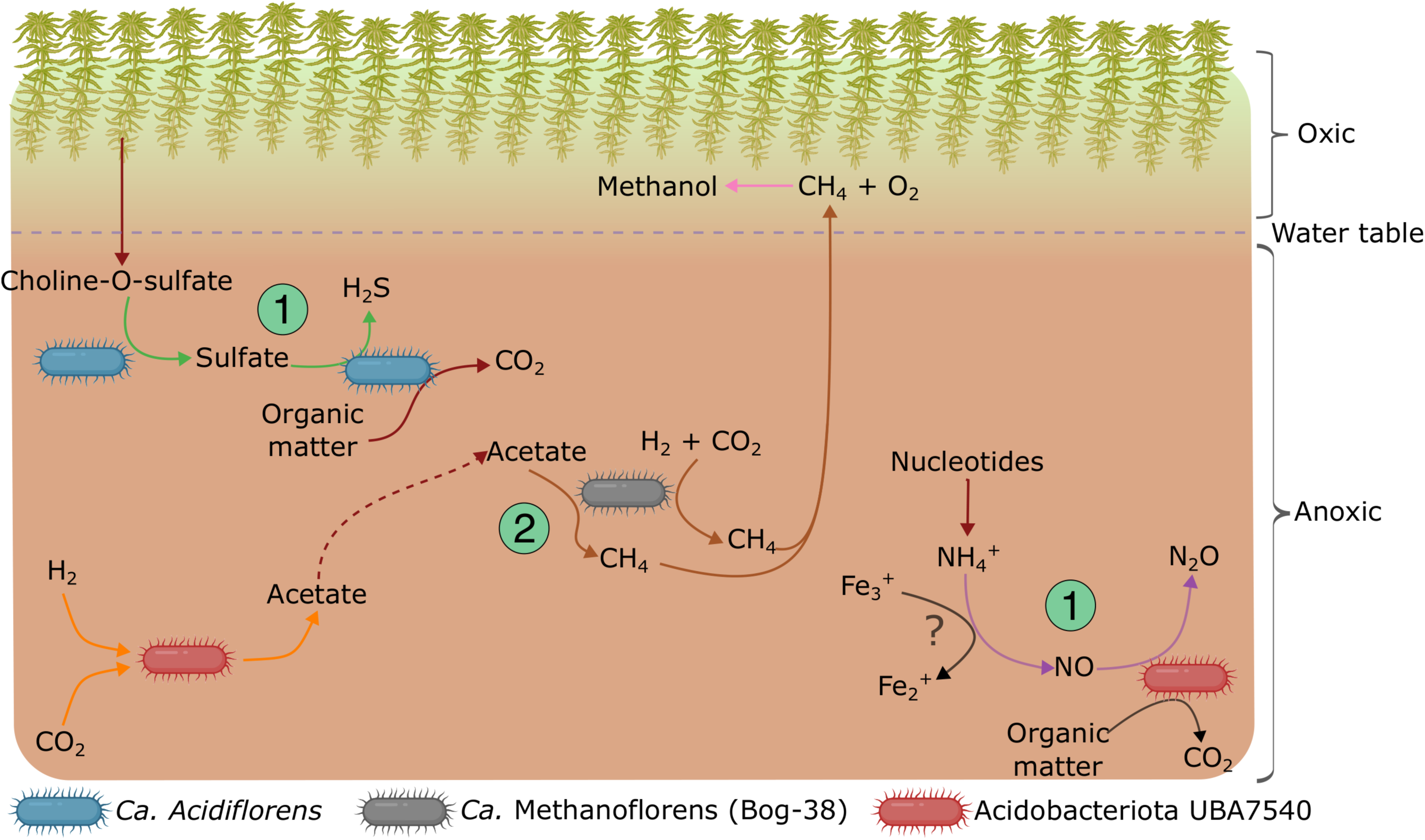
Conceptual model of terminal steps of organic matter degradation in peatlands. Simplified model of the main terminal electron-accepting pathways driving carbon turnover in SPRUCE peat based on the data shown in Figures 3 and 4. Arrow colors follows the same schema as in Figure 4A. Dashed lines show indirect interactions while question marks point to reactions that could hypothetically be taking place in SPRUCE peat based on literature and genomic potential. 1) Cleavage of organic matter (choline-O-sulfate and nucleotides) to obtain TEA to fuel anaerobic respiration. 2) Metabolic versatility of methanogens with the capability to perform both acetoclastic and hydrogenotrophic methanogenesis.

First, the wide metabolic versatility of abundant species, may allow them to adapt their activity without significantly changing their abundance. For example, *Ca.* Methanoflorens (Bog-38) species, exhibit a genomic repertoire capable of performing both acetoclastic and hydrogenotrophic methanogenesis. Thus, these versatile methanogens can take advantage of the available acetate, or if acetoclastic methanogenesis is not energetically favorable, use the CO_2_ and H_2_ produced from the oxidation of energetically rich substrates for hydrogenotrophic methanogenesis (**Figure 5**)(Horn *et al*., 2003). This metabolic versatility, in combination with the slow growth rate of methanogens (Thauer *et al*., 2008; Conrad, 2023), alongside the influence of temperature on microbial activity, may explain why biogeochemical data indicates methanogenic activity increases with warming, but community composition remains the same. A transition between methanogenic pathways with increasing temperature is indeed expected, based on previous isotopic measurements in the SPRUCE peat that indicated a shift from hydrogenotrophic to acetoclastic methanogenesis (Wilson *et al*., 2021). However, validation of such transition in response to climate change necessitates further experimental evidence from activity-based (metatranscriptomic) studies.

Second, the increased input of organic matter resulting from higher plant productivity at warmer temperatures also replenishes the pool of TEA available for anaerobic respiration, given the potential capability of peat microbes to obtain TEA from organic compounds (**Figure 5**). This mechanism could thus be facilitating the respiration of organic matter via alternative pathways to methanogenesis and thus, acting as a counterbalance to limit the growth and activity of methanogens by fostering the activity of other anaerobic groups (e.g., sulfate-reducers) and thus, the release of CO_2_. The genomic evidence reported here in support of this mechanism is further complemented by previous studies that showed that TEA limitation is alleviated at high temperatures (Tveit *et al*., 2015; Song *et al*., 2023). Experimental data will be required in the future to assess the existence and extent of this mechanism. For example, the activity of enzymes responsible for the cleavage of TEA from organic compounds, such choline-sulfatase (*betC*), would be expected to correlate with the release of CO_2_. Culture studies will also be needed to verify these physiological mechanisms.

To conclude, we have assessed the impact of warming on keystone functional guilds, providing evidence that microbial networks are resilient to warming. Therefore, the previously reported increase in CH_4_ and CO_2_ is likely due to changes in microbial activity rather than community changes. The uncovered taxonomic and metabolic novelty of microbes inhabiting peat soils along with our proposed conceptual model alter perceptions of anaerobic respiration pathways mediating terminal decomposition in peatlands and provide a mechanistic framework for improved predictions of the ratio of CO_2_:CH_4_ emissions and their likely response to climate drivers.

## Methods

### Study site

The Spruce and Peatland Responses Under Changing Environments (SPRUCE) is whole ecosystem warming experiment located in the ombrotrophic, acidic S1 Bog of the Marcell Experimental Forest, north of Grand Rapids, MN (47°30.4760N; 93°27.1620W). The outflow chemistry from the enclosures indicates that, in august 2016 and 2018, on average the pH was 3.42, total organic carbon was 85.83 mg C/L, total nitrogen was 1.71 mg N/L, total phosphorous was 0.30 mg P/L, and total sulphate was 0.67 mg/L, (see **Supplementary Information**). The S1 Bog is dominated by peat mosses of the genus *Sphagnum*, including *S. fallax* and *S. divinum* (previously *magellanicum*), and sparsely populated by Black spruce (*Picea mariana*) and larch (*Larix laricina).* Ericaceous shrubs such as *Rhododendron groenlandicum* and *Chamaedaphne calyculata* are commonly found throughout the peatland and accompanied by the herbaceous *Maianthemum trifolium*. In the absence of severe drought conditions, the water table fluctuates at a depth of between 0 and 30 cm of peat depth (Lin *et al*., 2014b; Hanson *et al*., 2020; Malhotra *et al*., 2020; Iversen *et al*., 2023; Petro *et al*., 2023). More information about the experiment can be found in **Supplementary Information**.

### DNA extraction and metagenome sequencing

Peat samples were collected in June of 2015 and 2016 and August of 2018 from the hollows of each of the 10 whole-ecosystem warming chambers with a serrated knife at the surface and a Russian corer at depth. The samples were separated into depth increments and six 0.35-g subsamples of homogenized peat from the 10-20 cm, 40-50 cm, 100-125 cm, and 150-175 cm depth increments were collected and extracted for DNA using the MoBio PowerSoil Pro Kit (QIAGEN) and resuspended in 50 μL of 10 mM Tris buffer. Illumina TruSeq metagenome libraries were generated by the JGI using their standard protocol. Metagenomes are publicly available on the JGI Genome Portal and NCBI Sequence Read Archive (SRA) (**Table S4**).

### Metabolomics

Metabolomics analysis was performed on biologically replicated wet peat samples collected in 2018 that were also used for metagenomic analysis. Briefly, peat samples were lyophilized, then extracted using a mixture of MeOH and sterile water. Samples were sonicated, filtered, freeze-dried, and stored at −80 °C before analysis by liquid chromatography–mass spectrometry (LC-MS/MS). Before analysis, samples were reconstituted in water:methanol for reverse phase (RP) and water:acetonitrile for hydrophilic interaction liquid chromatography (HILIC). Spectral data collection was carried out using a Thermo Scientific Orbitrap Exploris 480 mass spectrometer. Data analysis was conducted using the Compound Discoverer 3.3 software by Thermo Fisher Scientific, employing an untargeted metabolomics workflow (see **Supplementary Information** for further method details).

### Metagenome preprocessing, binning and annotations

Metagenomic reads were quality filtered using bbduk.sh v38.18 (qtrim=w,3 trimq=17 minlength=70 tbo tossjunk=t cardinalityout=t) (Bushnell B., 2015) and then, coverage was normalized using bbnorm.sh v38.18 (target=30 min=5 prefilter=t tossbadreads=t). Both original and normalized metagenomic reads were assembled independently with idba_ud v1.1.3 (--maxk 120) (Peng *et al*., 2012) and SPAdes.py v3.15.5 (-k 21,33,55,77,99,127 --meta --only-assembler) (Bankevich *et al*., 2012). Then, contigs longer than 1kb were used to recovery metagenome-assembled genomes (MAGs) with Maxbin v2.2.7 (Wu *et al*., 2016) and metaBAT2 v2.15 (Kang *et al*., 2019). MAGs were first dereplicated (within samples) with miga v1.3.8.3 (derep_wf --fast) (Rodriguez-R *et al*., 2018) yielding a total of 5,963 MAGs that were further dereplicated (between samples) using dRep v3.4.3 (-comp 50 -con 15 -sa 0.95 --S_algorithm fastANI) (Olm *et al*., 2017). As a result, the final dataset was composed of 697 unique genomospecies. CheckM v1.2.2 (lineage_wf) (Parks *et al*., 2015) was employed to assess MAG quality and GTDB-tk v2.1.0 (classify_wf) (Parks *et al*., 2020) to classify them taxonomically. MAG annotation was performed with DRAM using default parameters (Shaffer *et al*., 2020) and KEGG module completeness was evaluated using a custom R script (see **Table S5** for criteria used to define presence/absence of each metabolic pathway). The potential metabolic pathways for each MAG are available in **Table S6**. CoverM v0.6.1 (Woodcroft, 2021) was employed to calculate the truncated average sequencing depth at 80% (TAD80) for each MAG (-p bwa-mem --min-read-aligned-length 100 -- min-read-percent-identity 95 --min-read-aligned-percent 70 --exclude-supplementary -m trimmed_mean --trim-min 10 --trim-max 90). TAD80 was then normalized by genome equivalent values obtained from MicrobeCensus v1.1.0 (Nayfach & Pollard, 2015) to obtain relative abundances. MAGs analyzed in this manuscript were deposited in NCBI under BioProject PRJNA1084886.

### Phylogenomic tree

Proteins encoded on each MAG were first predicted using Prodigal v2.6.3 (-p meta) (Hyatt *et al*., 2010).Then, PhyloPhlAn v3.0.67 was used to build a phylogenomic tree based on the alignment of 400 universal gene markers (Asnicar *et al*., 2020)(--diversity high --fast). The tree was finally drawn and annotations added on R (R Core Team, 2023) using ggtree (Yu *et al*., 2017).

### Comparison to genomes retrieved from representative global soil types

The largest metagenomic dataset from northern peatlands published to date was obtained from Stordalen Mire in Sweden (Cronin *et al*., 2023). To assess how different or similar our metagenomes are to Stordalen Mire metagenomes, approaches at both the read level and the MAG level were performed. As for the read level, MASH v1.1 (Ondov *et al*., 2016) distances were calculated (sketch -s 10000 -r -m 2) and plotted on an Non-metric MultiDimensional Scaling (NMDS) plot on R using the vegan (Dixon, 2003) and ggplot packages (Valero-Mora, 2010). Regarding the MAG level, all MAGs publicly available from Stordalen Mire (Cronin *et al*., 2023) were retrieved and dereplicated using dRep, as explained above. Then, ANI between dereplicated Stordalen Mire and SPRUCE MAGs (1,645 and 697 MAGs, respectively), were calculated with fastANI v1.33 (Jain *et al*., 2018).

To compare SPRUCE genomes to those from a variety of representative soils around the world, metagenomes from upland forest (Hubbard Brook Experimental Forest)(Roco *et al*., 2019), Antarctic (Liu *et al*., 2021), tropical (Liu *et al*., 2021), grassland (Luo *et al*., 2014), agricultural soils (Orellana *et al*., 2018) as well as peat incubation samples from Ward reservation (Massachusetts, USA) (Reji & Zhang, 2022) were retrieved from the SRA (**Table S7**). MASH distances were calculated, as explained above, and 3D NMDS plots were drawn with the plotly R package (Sievert, 2020).

### Gene abundances

Genes and proteins were predicted from all contigs longer than 1 kb recovered from the 131 metagenomes using Prodigal v2.6.3 (-p meta). MMseqs2 v13.45111 (Steinegger & Söding, 2018) was employed to obtain a non-redundant gene database (easy-cluster -c 0.8 --cov-mode 0 --min-seq-id 0.95) which was then used by CoverM v0.6.1 to calculate gene sequencing depth (-p bwa-mem --min-read-percent-identity 95 --min-read-aligned-percent 50 --exclude-supplementary -m mean --min-covered-fraction 80 --contig-end-exclusion 20). The non-redundant gene ids were extracted from the protein file using FastA.filter.pl from the enveomics package (Rodriguez-R & Konstantinidis, 2016) to get the non-redundant protein database which was annotated with kofamscan (Aramaki *et al*., 2019), and the output filtered to keep only hits with evalue < 1e-15 and scores > 90% of the precomputed scores. Gene sequencing depth and annotation data was merged with Table.merge.pl and the table parsed with custom R scripts to get the KO sequencing depth per sample. Finally, KO relative abundances were obtained by dividing the KO sequencing depth by genome equivalents.

### Statistical analysis

When testing for the impact of warming and eCO_2_ on MAG relative abundance, only samples from 2016 and 2018 were used since warming was initiated in 2015, and eCO2 in 2016. Statistical analyses were done in R version 4.2.3 (R Core Team, 2023). Bray-Curtis and Unifrac beta diversity were calculated from the MAG relative abundance (distance(), phyloseq) (McMurdie & Holmes, 2013). The resulting matrices were used as the response variable of a distance based redundancy analysis (dbrda(), vegan), with the full model including sampling read and depth. Stepwise model selection was used to select the best fit model (ordiR2step(), vegan). Depth specific models were created to determine if warming and eCO2 influenced community composition. This resulted in a total of four models, and temperature was calculated from the average over the month of August during the year of collection at the probe closest to the depth fraction. For the samples at 20-30 cm of depth we used the temperature at 20 cm, at 40-50 cm we used the temperature at 40 cm, at 100-125 cm we used the temperature at 100 cm, and for 150-175 we used the temperature at 200 cm.

To target specific hypotheses, we fitted multiple linear regressions for the aggregated relative abundance of MAGs in 2016 and 2018 that had the metabolic potential for pathways of interest as the response variable (lmer, lme4)(De Boeck *et al*., 2011). Pathways of interest included homoacetogenesis via the Wood-Ljungdahl pathway, acetogenesis via fermentation, sulfate and sulfite reduction, and methylotrophic, acetoclastic, and hydrogenotrophic methanogenesis. Response variables included peat temperature and eCO_2_, and we included year as a random factor, the step function (lmerTest) was used to eliminate non-significant predictors (Kuznetsova *et al*., 2017). We also employed Maaslin2 (Mallick *et al*., 2021) to determine multivariable association between temperature and all MAG abundances (min_prevalence = 0) in 2018. We did not include 2016 since year-to-year effects could not be accounted for. Fixed effects included warming only. For both hypothesis-based models and Maaslin2, each depth was process separately, and temperature was calculated as for the distance-based redundancy analysis.

LC-MS/MS feature peak abundance values were normalized through mean normalization. Distance matrices were calculated via Bray-Curtis dissimilarity and used for permutational multivariate ANOVA (PERMANOVA) in the vegan package, with visualization in ggplot2. Log2 fold change (L2FC) values were computed for each identified feature to identify meaningful differences in compound expression across different depth. T-tests were performed to determine statistical significance between depths. The Benjamini-Hochberg procedure was used to control the false discovery rate, which provided adjusted p-values (Padj) to account for multiple testing (Benjamini & Hochberg, 1995). Adjustments were made separately for RP and HILIC datasets. Compounds were then filtered based on associated Padj. values <0.05, categorizing them as upregulated in the sample when L2FC > 0, or downregulated in the sample when L2FC < 0.

International chemical identifier keys (InChIKeys), which provide standardized, condensed text representations of chemical compounds, were extracted. These InChIKeys were utilized to classify compounds into standard chemical taxonomy categories using ClassyFire (Djoumbou Feunang *et al*., 2016). Integration of metabolomic and metagenomic data (gene abundances) was performed with the DIABLO workflow of the R package mixOmics (Singh *et al*., 2017) using clr-transformed data (clr, compositions R package)(van den Boogaart & Tolosana-Delgado, 2008). For the correlation analysis, the function circosPlot was used and a minimum correlation score of 0.8 or −0.8 was set to call positively and negatively correlated variables.

