## Supplementary Information for "Northern peatland microbial networks exhibit resilience to warming and acquire electron acceptor from soil organic matter"

*equal first authors

**Supplementary methods**

*SPRUCE experiment*

The experiment consists of 17 open-top chambers that control the peat and air temperature (ambient, +0, +2.25, +4.5, +6.75 and +9°C) as well as atmospheric CO_2_ concentration (ambient and 900 ppm). In June 2014, an array of heating rods 3-m in depth were established and began warming the belowground peat. A subsurface corral hydrologically isolates each enclosure but allow natural lateral water flow. In August 2015, aboveground air warming was initiated. The air within the enclosure is heated by a propane heater outside the chamber and reintroduced to achieve desired temperature levels, with monitoring conducted at the target control point located at +2 meters in the center of the plot. In 2016, CO_2_ additions began. The pure CO_2_ is vaporized and warmed before being distributed and homogenized within the enclosure (Hanson *et al.*, 2017). Environmental variables such as temperature, precipitation and gas emissions are constantly monitored. The readers are referred to the official webpage for further information on the experiment and a comprehensive list of project measurements (<https://mnspruce.ornl.gov>).

*Calculation of outflow chemistry from SPRUCE datasets*

We used outflow water autosampled from the experimental enclosures as reported in Sebestyen et al., 2021. We calculated the mean pH, sulfate, total nitrogen, total phosphorous, and total organic carbon from all measurements taken in the month of August.

*Metabolomics*

Metabolomics analysis was performed on biologically replicated wet peat samples collected in 2018 that were also used for metagenomic analysis. To dry samples and ensure uniform starting weight for extraction, peat samples were first lyophilized using a Labconco FreeZone, Benchtop freeze dryer for 48 hr. The freeze-dried peat samples (0.2 g) were extracted by adding 20 mL of an 80:20 solution of MeOH: sterile MilliQ water. Samples were briefly vortexed and sonicated in a water bath for 2 hr at 20 °C (FisherBrand CPX3800). The supernatant was filtered through a 0.45 um filter to remove cellular debris and plant material. Of this extract, 7 mL was transferred to two glass autosampler vials (3.5 mL each), dried in a vacuum centrifuge (Eppendorf Vacufuge plus), and stored at −80 °C. Prior to CL-MS/MS analysis, samples were reconstituted in 80:20 water: methanol for reverse phase (RP), and 50:50 water: acetonitrile for hydrophilic interaction liquid chromatography (HILIC).

### *Liquid chromatography-tandem mass spectrometry (LC-MS/MS)*

Liquid chromatography was conducted using a Thermo Scientific Vanquish Duo ultra-high performance liquid chromatography system (UHPLC). For reverse phase (RP) separation, extracts were separated on a Waters ACQUITY HSS T3 C18 column, while hydrophilic interaction liquid chromatography (HILIC) separation utilized a Waters ACQUITY BEH amide column. Samples were injected into the column with a volume of 1 microliter. The elution conditions were as follows: for RP, a gradient from 99% mobile phase A (0.1% formic acid in H2O) to 95% mobile phase B (0.1% formic acid in methanol) was employed for 16 minutes. For HILIC, the gradient went from 99% mobile phase A (0.1% formic acid, 10 mM ammonium acetate, 90% acetonitrile, 10% H2O) to 95% mobile phase B (0.1% formic acid, 10 mM ammonium acetate, 50% acetonitrile, 50% H2O). Both columns were operated at a temperature of 45 °C, with a flow rate of 300 microliters/minute.

Spectral data collection was carried out using a Thermo Scientific Orbitrap Exploris 480 mass spectrometer. For RP, a spray voltage of 3500 V was applied in positive mode, while for HILIC, 2500 V was used in negative mode with the H-ESI source. The ion transfer tube and vaporizer temperature were maintained at 350 °C. Compound fragmentation was achieved through data-dependent MS/MS with HCD collision energies set at 20, 40, and 80.

*Metabolite processing and annotation*

Data analysis was conducted using the Compound Discoverer 3.3 software by Thermo Fisher Scientific, employing an untargeted metabolomics workflow. The initial steps of the analysis involved spectral alignment and peak picking. Putative elemental compositions of unknown compounds were predicted using the exact mass, isotopic pattern, fine isotopic pattern, and MS/MS data using the built in HighChem Fragmentation Library of reference fragmentation mechanisms. Metabolite annotation was performed using an in-house database built from 1200 reference standards, spectral libraries and compound databases. First, fragmentation scans, retention time and ion mass of unknown compounds were compared with those in the in-house database. Second, fragmentation scans (MS2) searches in mzCloud were performed, which is a curated database of MSn spectra containing more than 9 million spectra and 20000 compounds. Third, predicted compositions were obtained based on mass error, matched isotopes, missing number of matched fragments, spectral similarity score (calculated by matching theoretical and measured isotope pattern), matched intensity percentage of the theoretical pattern, the relevant portion of MS, and the MS/MS scan. The mass tolerance used for estimating predicted composition was 5 ppm. Finally, annotation was complemented by searching MS1 scans on different online databases with ChemSpider (using either the exact mass or the predicted formula). To enhance annotation coverage and add compound classes, SIRIUS, CSI:FingerID, and CANOPUS were used (Kai et al., 2019). Compounds chemical taxonomy assignment based on chemical structure was assigned using ClassyFire (Djoumbou Feunang et al., 2016). Furthermore, metabolite annotation was further enhanced using CMM-RT which employs machine learning, artificial intelligence, and neural networks for more accurate predictions of Retention Times (RTs) (Garcı´a et al., 2022). Compounds level of annotation was assigned according to the Metabolomics Standards Initiative (Sumner et al., 2007) as follows: Level 1: compounds with exact match to a standard reference compound in our in-house library, Level 2: compounds with full match to online spectral databases using mzCloud database (based on MS2 spectra matching), Level 2.1 for compounds with a full match to databases such as ChemSpider and/or CMMRT, relying on mass, molecular formula, and/or retention time, combined with annotations from SIRIUS using MS2 fragmentation patterns, Level 2.2 applied to compounds with a full match to databases (ex. ChemSpider ...) and CMMRT based on mass, molecular formula, and retention time, Level 2.3 encompassed compounds matching any single annotation source, Level 3.1: for matches based on molecular formulas generated through Predicted Compositions node and/or CMMRT, combined with annotations from SIRIUS using MS2 fragmentation patterns, Level 3.2 indicated matches based on molecular formulas generated through Predicted Compositions in conjunction with CMMRT. Finally, Level 3.3 represented matches based on chemical formulas from any annotation source. Following metabolite annotation, poorly annotated features were eliminated by performing manual QC on the features. QC correction was applied in Compound Discoverer using a linear regression model that retains features with a QC area RSD < 30% and was limited to a maximum correction of < 25%. Mass spectra and chromatography from duplicate annotations (for a single compound) were manually inspected for spectral quality, peak shape, and retention time following data processing with Compound Discoverer to assess if one of the multiple annotations was correct. When a single confident annotation was identified, all other duplicate feature annotations were removed. Further annotations were inspected for removal of in-source fragments.

**Supplementary results**

*Summary of the metabolic profile of the peat in 2018*

Through untargeted metabolomics, we detected a total of 5261 features across different depths, which we annotated and identified to varying degrees using Metabolomics Standards Initiative (MSI) criteria (Sumner et al., 2007). We confidently identified 136 features (2.6%) as Level 1 annotations, providing the highest level of certainty. Additionally, we putatively annotated 527 (10%) as Level 2 and tentatively characterized 63 features (1.2%) as Level 2.1, 546 features (10.4%) as Level 2.2, and 638 features (12.1%) as Level 2.3. In total, we assigned chemical names and molecular formulas to 1,910 features, representing a substantial portion of our dataset (more details are provided in the methods section). This extensive annotation effort is essential for converting complex metabolomics data into practical biological insights. Our annotation efforts also proposed molecular formulas for 488 features (9.3%) as Level 3.1 annotations, 1090 features (20.7%) as Level 3.2, and 601 features (11.4%) as Level 3.3 annotations, resulting in 2,179 features with at least a formula. Furthermore, 692 features (13.2%) had multiple potential annotations emphasizing the complexity of the metabolome, while 9.1% remained unannotated, suggesting the presence of novel metabolites.

Lipids and lipid-like molecules were the most commonly identified molecules (335 compounds), and the category was dominated by fatty acyls, prenol lipids, and steroids and steroid derivatives. Organic acids and derivatives were the second most common identified compound (244 compounds, with 214 classified as carboxylic acids and derivatives), followed closely by organoheterocyclic compounds (226 compounds). The metabolites, like the metagenome assembled genomes, were strongly depth stratified (**Figure S4**). This suggests depth is a key factor influencing metabolic diversity in the peat system. Surface depths (10-20cm, 40-50cm) harbored the most distinct compositions compared to deep layers (100-125cm, 150-175cm). Metabolic diversity was highest in the surface layers, with an average Shannon alpha diversity of 6.11, 6.08, and 6.05 at 10-20, 40-50, and 100-125 cm respectively. The least diverse layer was the very deep peat, the 125-150 cm depth increment, with a Shannon diversity index of 5.82.
