## Supplementary Figures for "Northern peatland microbial networks exhibit resilience to warming and acquire electron acceptor from soil organic matter"

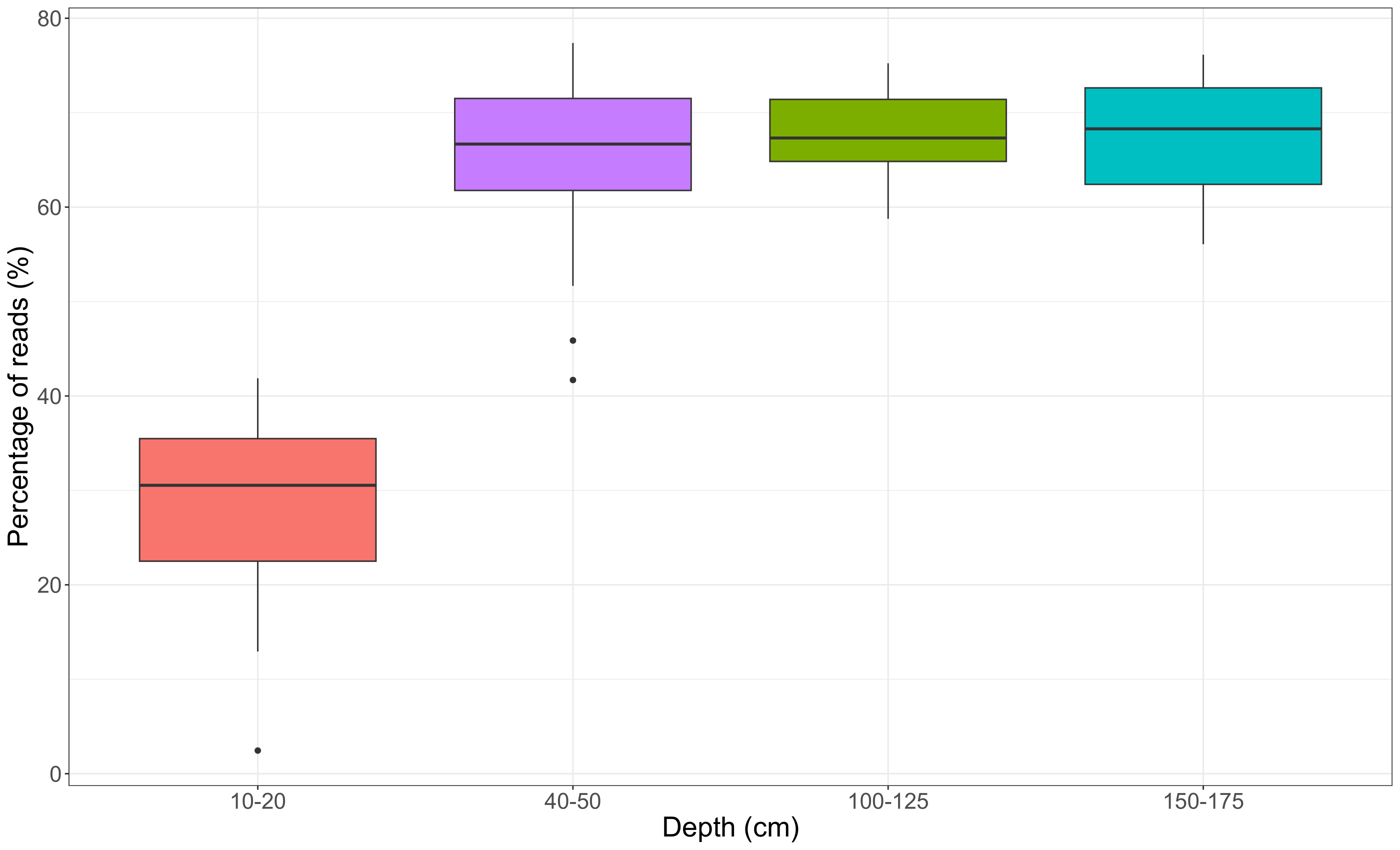

**Supplementary Figure 1.** Percentage of metagenomic reads recruited against the MAG dataset per depth. Note the lower read recruitment in the surface layer (10-20 cm) compared to the other three deeper layers.

See Figure S2.html file on an internet explorer

**Supplementary Figure 2.** 3D NMDS plot based on MASH distances of raw reads comparing different types of soil metagenomes (agricultural, antartic, grassland, peat (3 sites: SPRUCE, Stordalen Mire and Ward reservation), tropical forest, upland forest.

See Figure S3.html file on an internet explorer

**Supplementary Figure 3.** 3D NMDS plot based on MASH distances of raw reads comparing SPRUCE bog to the bog, palsa and fen in Stordalen Mire.

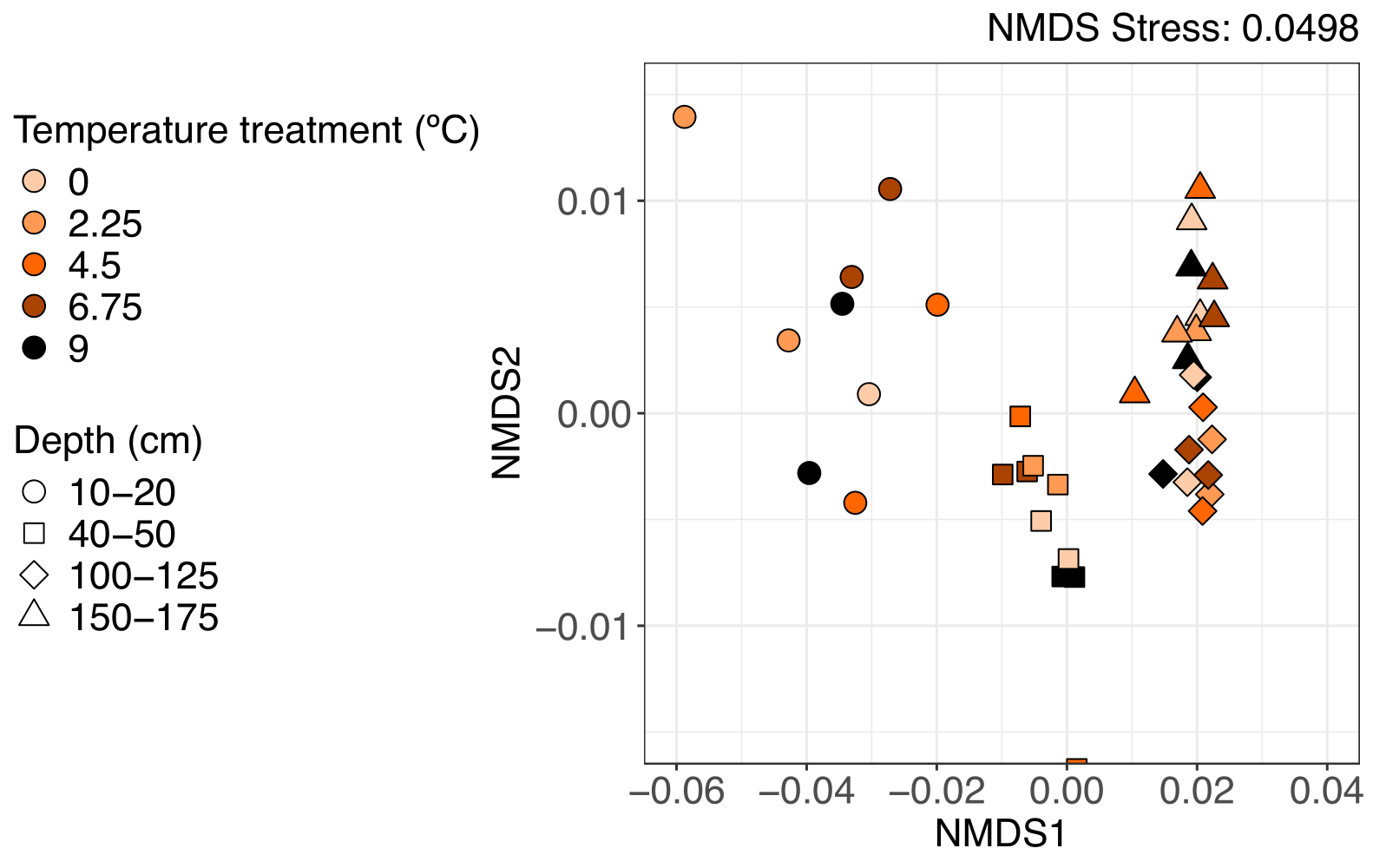

**Supplementary Figure 4.** NMDS plot based on metabolite composition and abundance in peat samples collected in August 2018. Shapes indicate the four depths (10-20, 40-50, 100-125 and 150-175 cm) while colors show the temperature treatment

**Supplementary Figure 5.** Average relative abundance (bubble plot on the left side) and metabolic potential (heatmap on the right side) for the top 50 most abundant MAGs in SPRUCE peat (rows). MAGs are sorted by z-score, the most prevalent at the surface layer at shown at the top while the most abundant at the deepest layer are shown at the bottom. Note that the potential for aerobic respiration and fermentation predominates at the surface layer, decreasing in importance with depth. On the contrary, anaerobic respiration processes (sulfate/sulfite reduction, denitrification and methanogenesis).
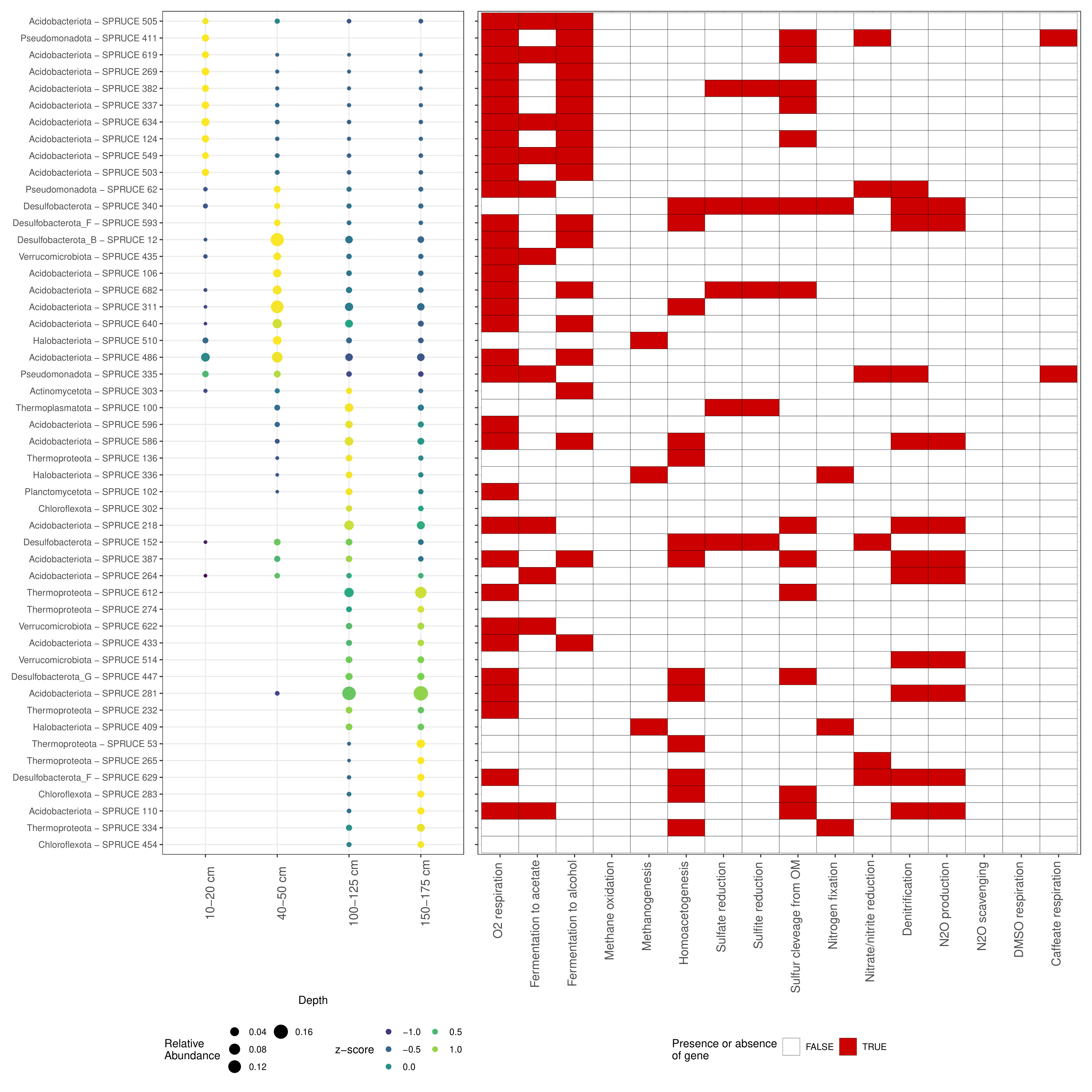

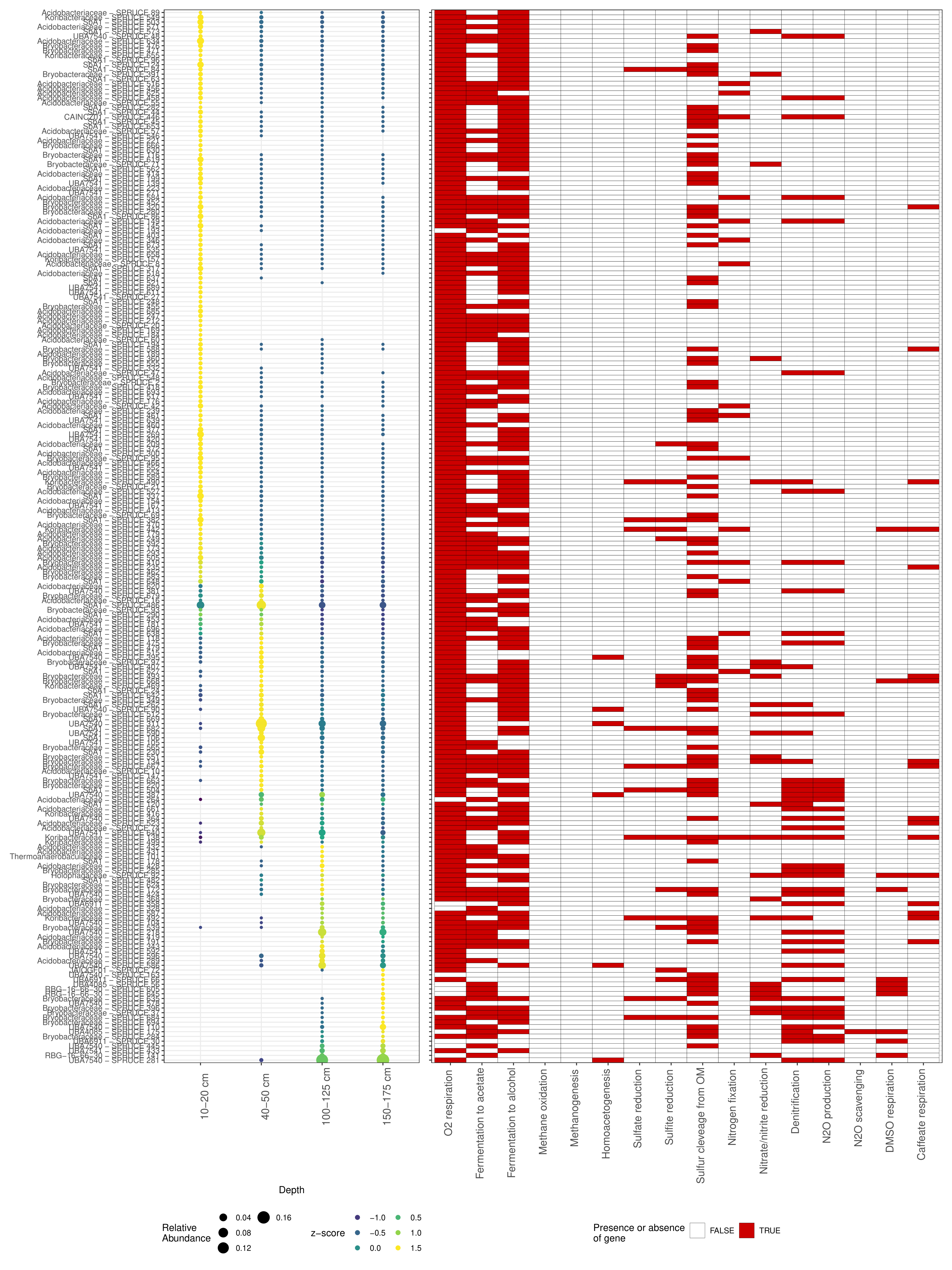

**Supplementary Figure 6.** Average relative abundance (bubble plot on the left side) and metabolic potential (heatmap on the right side) for the *Acidobacteriota* MAGs in SPRUCE peat (rows). MAGs are sorted by z-score, the most prevalent at the surface layer at shown at the top while the most abundant at the deepest layer are shown at the bottom.

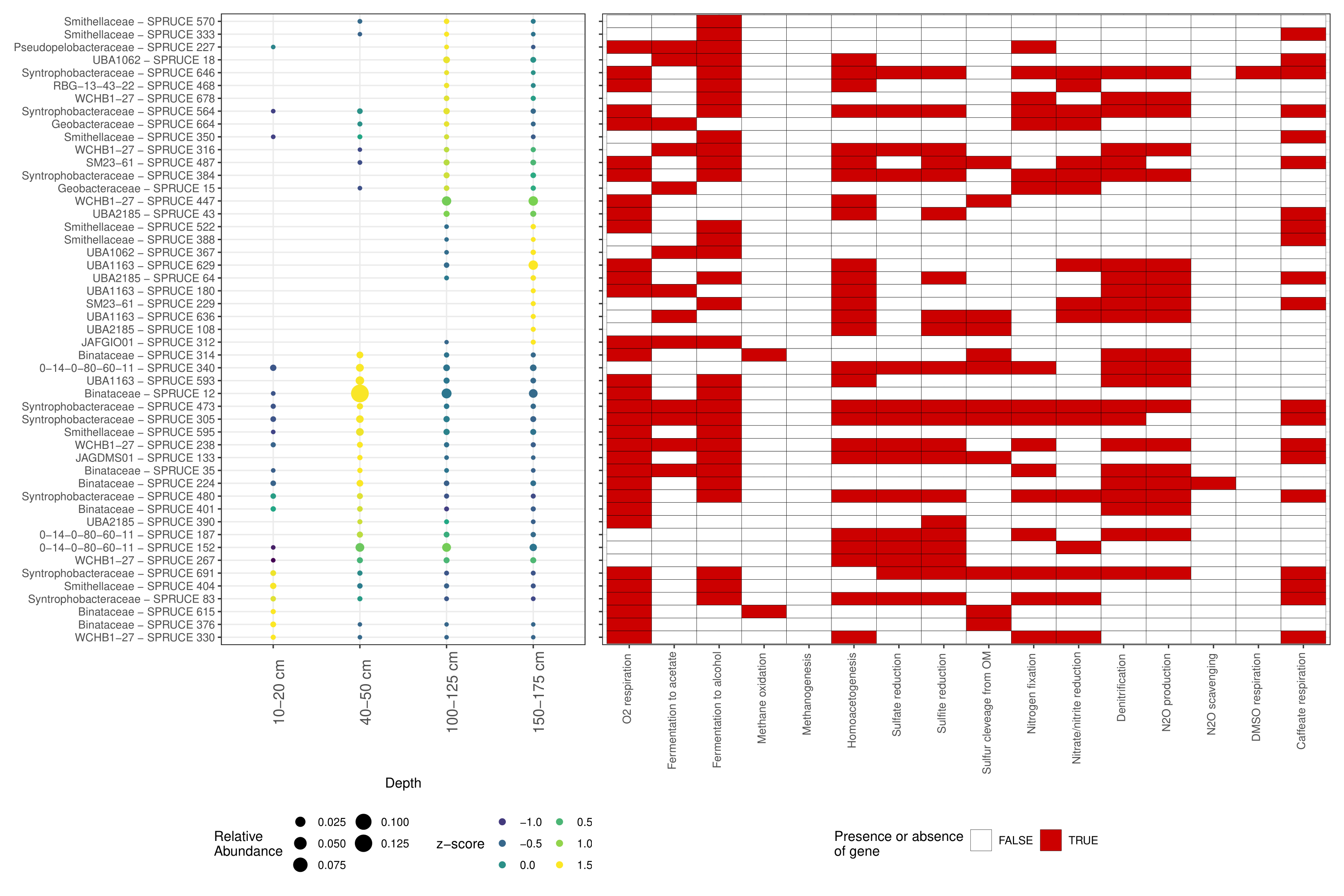

**Supplementary Figure 7.** Average relative abundance (bubble plot on the left side) and metabolic potential (heatmap on the right side) for the *Desulfobacterota* MAGs in SPRUCE peat (rows). MAGs are sorted by z-score, the most prevalent at the surface layer at shown at the top while the most abundant at the deepest layer are shown at the bottom.

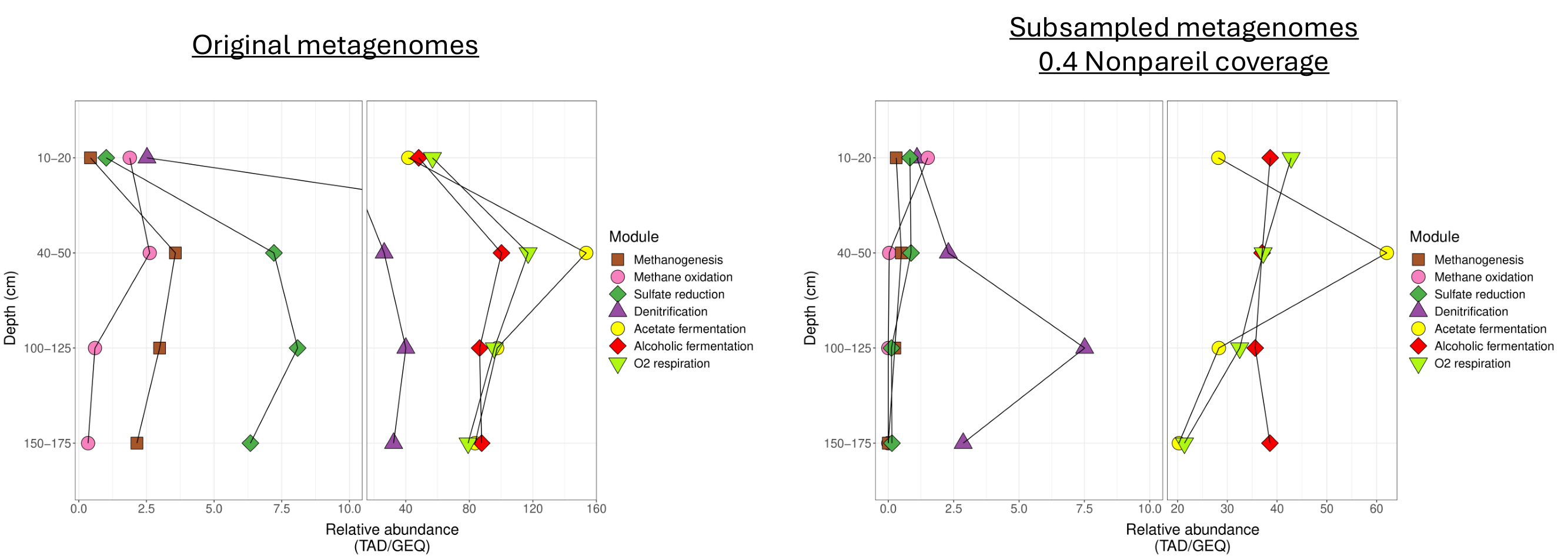

**Supplementary Figure 8.** Average relative abundance (TAD80/GEQ) of genes involved in central metabolic pathways in the SPRUCE peat calculated by read recruitment against genes using the original metagenomes (left side) and the Nonpareil coverage normalized metagenomes (right side). Note that continuous decay in oxygen respiration potential with depth on the Nonpareil coverage normalized metagenomes (right side).

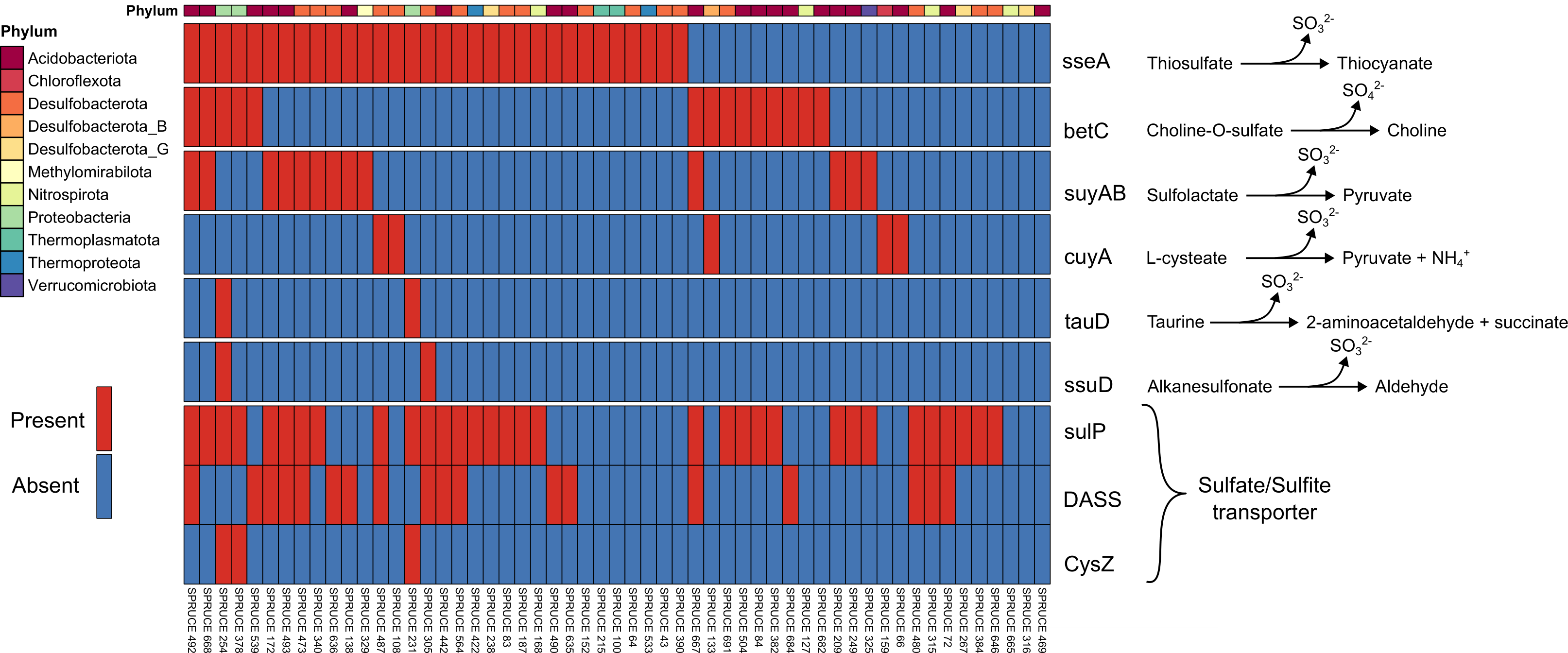

**Supplementary Figure 9.** Heatmap of the presence (red) and absence (blue) of genes involved in the cleavage of organic sulfur compounds and the consequent release of sulfate or sulfite (rows) in *dsrA* encoding MAGs (columns). The color bar at the top indicates the phylum each MAG belongs to. Note that most MAGs encode at least one gene involved in the cleavage of organic sulfur compounds.

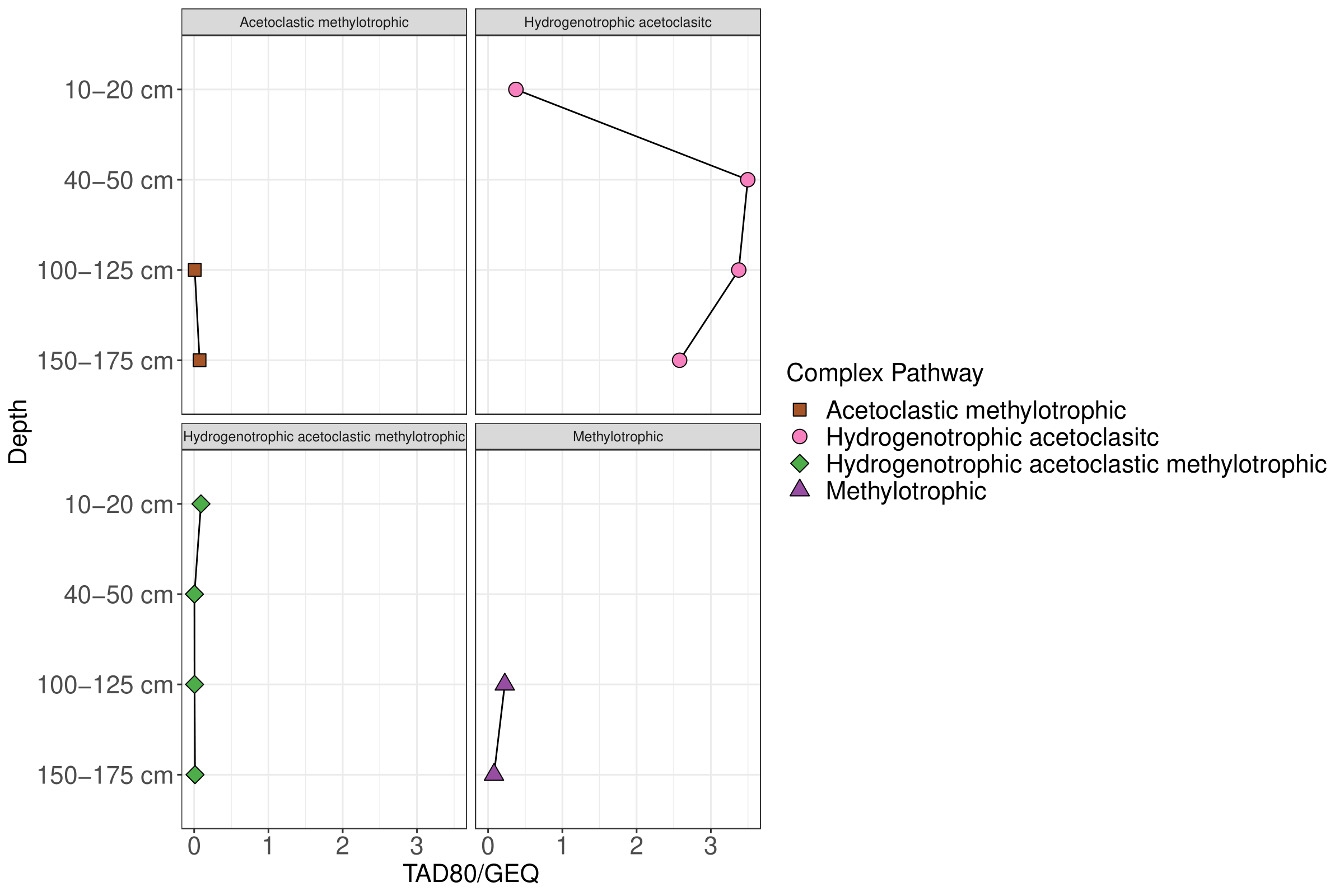

**Supplementary Figure 10.** Average relative abundance (x-axis, TAD80/GEQ) across the peat vertical profile (y-axis) of MAGs encoding methanogenic pathways. The relative abundance of MAGs was aggregated by metabolic potential (i.e., the abundance of MAGs with the same metabolic potential were summed up). Panels with more than one methanogenic pathway in the panel title show MAGs that encode multiple pathways in the same genome. Note that the potential for hydrogenotrophic and acetoclastic predominates at all depths but the potential for methylotrophic methanogenesis increases in the deepest layers (100-125 and 150-175 cm).

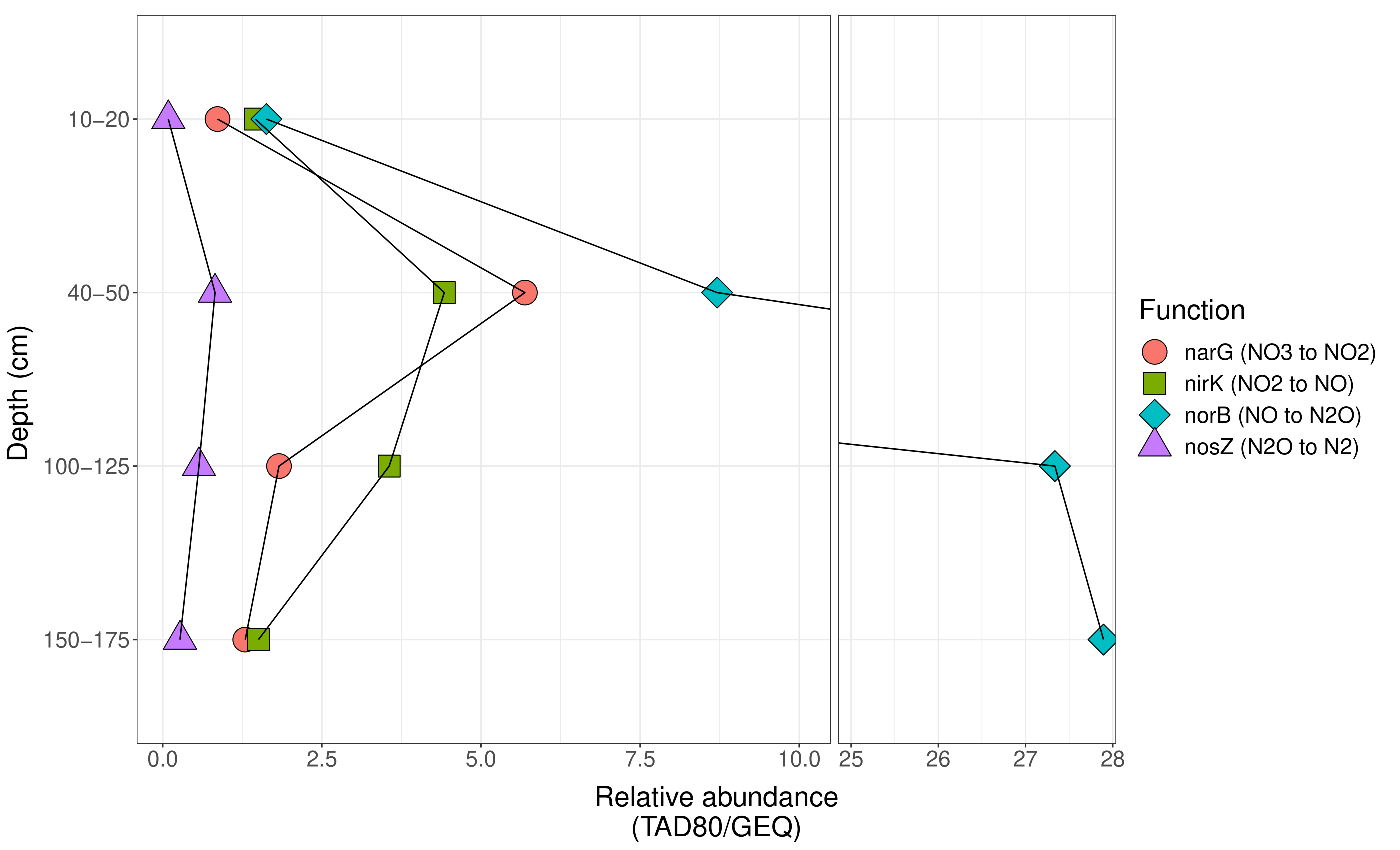

**Supplementary Figure 11.** Average aggregated relative abundance (TAD80/GEQ) of the MAGs encoding KOs involved in denitrification (narG = K00370; nirK = K00368; norB = K04561; nosZ = K00376). Note the predominance of *norB* at all depths but mainly at the 100-125 and 150-175 cm depths.

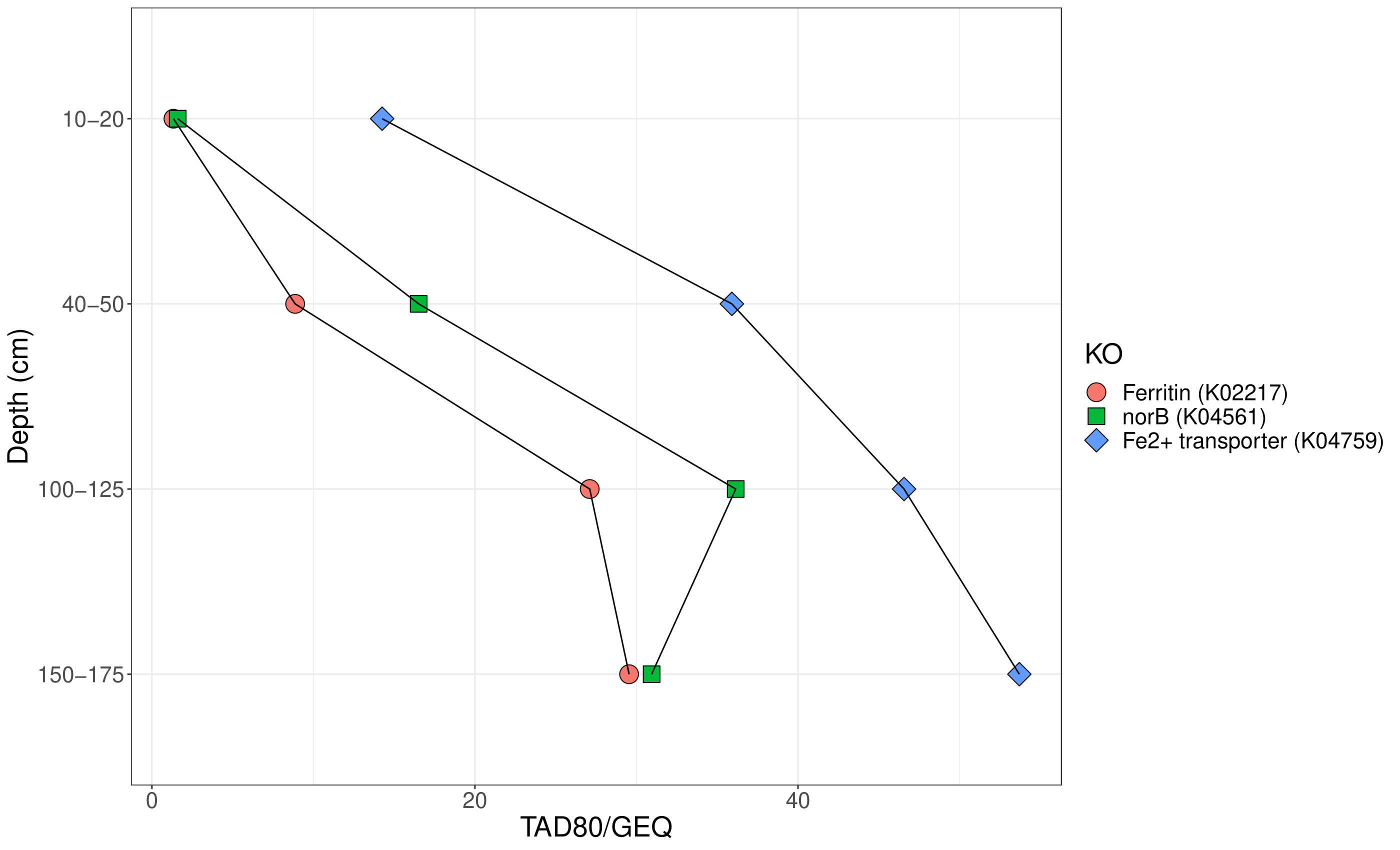

**Supplementary Figure 12.** Average relative abundance (TAD80/GEQ) of two KOs (K02217 and K04759) that are correlated with *norB* and are involved in the iron cycle. Abundance data was obtained by read recruitment against genes predicted from contigs.
